# Opposing effects of rewarding and aversive stimuli on D1 and D2 types of dopamine-sensitive neurons in the central amygdala

**DOI:** 10.1101/2024.11.07.622413

**Authors:** Łukasz Bijoch, Paweł Szczypkowski, Justyna Wiśniewska, Monika Pawłowska, Karolina Hajdukiewicz, Radek Łapkiewicz, Anna Beroun

**Author notes:** Corresponding author: Anna Beroun. Lead contact: Anna Beroun.

## Abstract

Dopamine-sensitive neurons are organized in two classes of cells, expressing D1- or D2- types of dopamine receptors, and are often mediating opposing aspects of reward-oriented behaviors. Here, we focused on dopamine-sensitive neurons in the central amygdala – a brain structure critically involved in processing emotion-related stimuli. We discovered that both dopamine receptor types are present in the central medial nucleus, while the lateral part is populated predominantly with DRD2 cells. Exposing mice to rewarding and aversive stimuli we studied DRD1 and DRD2 cells activity using *in vivo* two-photon calcium imaging in the CeM. We showed that cocaine and sugar predominantly increase the activity of DRD1(+) neurons and decrease DRD2(+) cells. Repeated exposure to cocaine, however, had the opposite effect on spontaneous excitatory synaptic transmission in the CeM than exposure to sugar. Quinine, an aversive stimulus, primarily engaged DRD2(+) neurons, activating predominantly those cells that were previously inhibited by sugar exposure. Our results show that though DRD1 and DRD2 populations are differentially engaged and regulated by appetitive/aversive stimuli, both participate in sugar, cocaine, and quinine processing.

## Introduction

The brain’s dopaminergic system is critically involved in reward and fear learning, as well as, responses to novel stimuli, serving as an indicator of novelty. This system also holds strong clinical importance, as dopaminergic dysregulation is implicated in various psychotic disorders such as addiction, with drugs of abuse directly or indirectly affecting its balance ^1,2^. Consequently, much effort has been made to characterize dopamine function in the mammalian brain. Though dopamine is released from two predominant brain areas: the substantia nigra (SN) and the ventral tegmental area (VTA), dopamine-sensitive neurons are widely distributed throughout the brain where they play a fundamental role in enabling motivation and reinforced learning, affect decision-making and attention, participate in various forms of learning and plasticity mechanisms to name just a few. The universally acknowledged model divides them into D1 and D2- types of cells, best described in the striatum where they reside in non-overlapping cellular populations. In general, D1-type cells express dopamine receptors D1 (DRD1) and D5 that bind to stimulatory G protein and promote cAMP production. Conversely, D2-type cells express dopamine receptors D2 (DRD2), D3, and D4 that decrease adenylate cyclase activity through inhibitory G protein binding. Thus, as dopamine elicits fundamentally different responses, depending on their type, cells expressing DRD1 or DRD2 are believed to opposingly regulate brain activity and participate in various aspects of reward-related behaviors. This opposing regulation has been particularly well-studied in the nucleus accumbens (NAC) – a key brain structure involved in the execution of motivated behaviors ^3–5^, where medium spiny projection neurons (MSN) expressing DRD1 or DRD2 respond differently to natural rewards (e.g. sucrose) and drugs of abuse (e.g. cocaine and morphine) ^6–8^. More recently, it became apparent that functionally and genetically distinct populations of the NAC DRD1 MSNs drive various aspects of reward- and addiction-related behaviors ^9^.

Although the striatum is crucial for addiction development, recent studies show that the central amygdala (CeA) is critically involved in reward processing and drug-related memory formation. Its function is to assign valence to stimuli and thus mediates appetitive and aversive memory formation that later drives behavior ^10–15^. The central amygdala with its subnuclei: medial, lateral, and capsular (CeM, CeL, and CeC, respectively) is functionally and genetically immensely heterogeneous, and its various distinct neuronal populations are recruited by appetitive and aversive stimuli ^11^. Unlike the basolateral nucleus of the amygdala (BLA), the CeA is highly populated with dopamine-sensitive neurons, though their exact function in response to rewarding stimuli remains largely unknown.

In this study, we investigated the involvement of dopamine-sensitive neurons in the CeA, in response to appetitive (cocaine and sugar) and aversive (quinine) stimuli. With an infrared camera, we recorded mouse facial expressions triggered by cocaine, sugar, and quinine administration. Two-photon in vivo imaging showed, that both rewards: cocaine and sugar predominantly activate DRD1 cells, while DRD2 cells inhibit their activity after cocaine and sugar exposure. In contrast, more DRD2 cells increase activity after an aversive stimulus. Looking at synaptic plasticity correlates (frequency of synaptic miniature events) after repeated rewards exposures, we observed opposing effects of cocaine and sugar on DRD1 and DRD2 cells, although chemogenetic manipulation of either DRD1 or DRD2 cell populations did not influence cocaine-conditioned place preference. Our results show that though DRD1 and DRD2 populations are differentially engaged by appetitive/aversive stimuli, both participate in sugar, cocaine, and quinine processing.

## Results

### DRD1 and DRD2 are expressed in non-overlapping populations in subnuclei of the CeA

The CeA, even though often described as a single entity, is a heterogeneous structure composed of nuclei with different functions and cell types ^11,13^. To study the expression patterns of dopamine receptors in its two major parts, CeM and CeL subnuclei, we examined the expression of reporter proteins TdTomato and EGFP in a transgenic mouse line: DRD1-TdTomato/DRD2- GFP ^16–18^.

We found that 44.59% of all cells in the CeL and 38.08% in the CeM were dopamine-sensitive. However, the composition of these cells varied between these nuclei: in the CeM DRD1(+) and DRD2(+) cells formed sparse populations, whereas in the CeL, almost all dopamine-sensitive neurons were DRD2(+) (Fig. 1). Cells co-expressing both these receptors were only a minor portion in both nuclei. This distinct composition pattern of CeM and CeL was also observed by labeling Cre protein in other transgenic mouse strains: DRD1-Cre and DRD2-Cre mice ^19,20^, (Fig. Supp1 F).

**Fig. 1:**
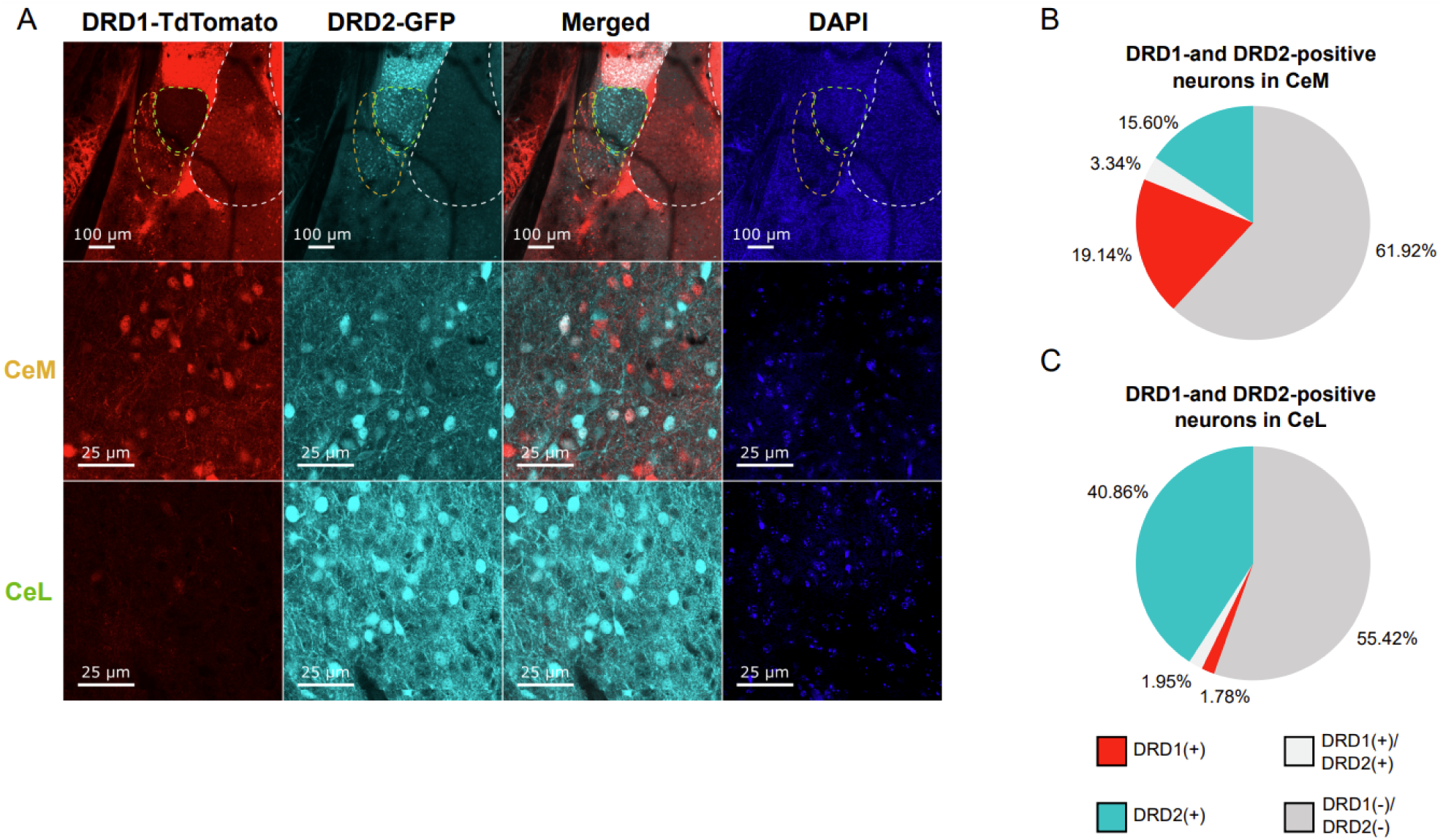
Different pattern of DRD1(+) and DRD2(+) cells in subnuclei of the central amygdala. (**A**) Exemplary images of the amygdala region from a DRD1-TdTomato (red) and DRD2-GFP-expressing cells (cyan) with DAPI-labeled nuclei (blue). Dashed lines represent subdivisions of the amygdala: white for the basolateral amygdala, green for the lateral nucleus of the central amygdala (CeL), and yellow for the medial nucleus of the central amygdala (CeM). The second row panels are images of DRD1- and DRD2-positive cells in CeM and the lowest row panels show cells from CeL. (**B, C**) Graphs representing percentages of DRD1(+), red, and DRD2(+), cyan, cells in CeM (**B**), and CeL nuclei (**C**). Graphs represent the average percentages of DRD1(+) and DRD2(+) cells in all the cells of CeM and CeL. For CeM N = 14(42) and for CeL N = 7(24), where N = number of animals (number of analyzed images).

A well-known marker of the CeA is protein kinase C delta (PKCδ), which is mostly limited to the CeL ^21^. Labeling this protein in slices obtained from D1-TdTomato/D2-GFP mice showed that 34.83% of PKCδ (+) in the CeM and 41.85% in the CeL were DRD2(+). Interestingly, only 2.41% of PKCδ (+) in the CeM were DRD1(+) (Fig. Supp. 1 A-E).

The dopaminergic system may also indirectly affect the CeA, as dopamine can bind to dopamine-sensitive neurons projecting to the CeA. To investigate this, we retrogradely labeled DRD1(+) and DRD2(+) neurons in DRD1-Cre and DRD2-Cre mice through stereotactic injections into the CeA (using retrograde pAAV-hSyn-DIO-EGFP or retrograde hSyn-DIO-mCherry). Afterward, we imaged samples, and using the semi-automatic software for structure annotation, we analyzed the brain for the presence of DRD1(+) or DRD2(+) cells. We found that the majority of dopamine-sensitive cells projecting to the CeA were DRD1(+), while DRD2(+) projected only from caudoputamen (CP) or the CeA, specifically from the CeC (Fig. Supp. 1 G, H). Conversely, DRD1(+) neurons projecting to the CeA were found in various parts of layer V of the cortex (determined as somatomotor, primary somatosensory, supplemental somatosensory, prelimbic, infralimbic, retrosplenial, agranular insular, gustatory, and visceral areas). Additionally, we also identified them in CP and zona incerta (ZI).

### Mouse behavior and facial expressions triggered by cocaine, sugar, and quinine exposure during *in vivo* two-photon imaging

Given that DRD1 and DRD2 cells forming non-overlapping populations are found in the CeM and not the CeL nucleus, we focused our attention on the CeM to study cell activity following cocaine intoxication, and sugar vs. quinine exposures.

We established a protocol for imaging the activity of CeM neurons, preceded by the gradual habituation of mice to head fixation (Fig. 2 A). To study the effects of reward exposures, we began the protocol by evaluating the effects of cocaine administration. Directly before each imaging session, once per day, mice received subcutaneous injections of either saline or cocaine solution (20 mg/kg body weight). Mice received saline on the first day, followed by two days of cocaine exposure, then saline, and a final cocaine injection. On the last imaging day, mice received droplets of liquid through a lickport. Each mouse received several droplets of 7.5% sucrose solution. Afterward, we administered an aversive stimulus - droplets of 0.1 mM quinine solution (Fig. 2 A).

**Fig. 2:**
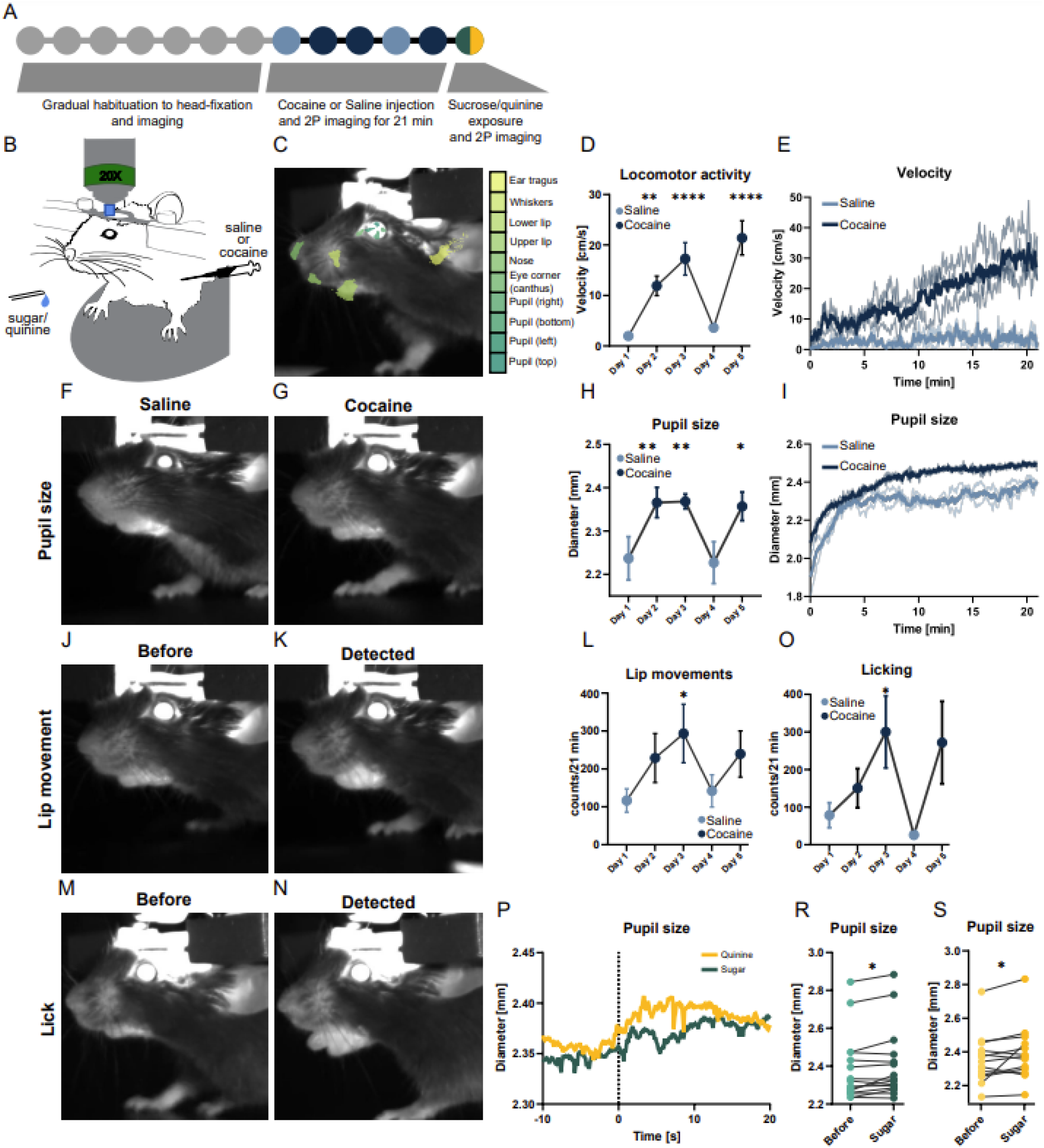
Facial expressions and locomotor activity of head-fixed mice exposed to cocaine, sugar, and quinine. (A) Experiment design. After the habituation period, mice were imaged under a two-photon microscope. Mice received once per-day subcutaneous injections of saline or 20 mg/kg body-weight cocaine solution. On the last day of the experiment, mice received droplets of 7.5% sucrose solution and 0.1 mM quinine solution as the last droplet. (B) Schematic representation of the mouse under the two-photon microscope. Head-fixed mouse stands on a rotating disc. (**C**) Facial features tracked with DeepLabCut software. Colored areas represent the positions of points detected by the software. (**D**) Graph representing the locomotor activity of a mouse on a running disc during cocaine and saline sessions. Data is presented as averages for all mice ± SEM. E) Averaged velocity of mice running on a rotating disc during saline (light blue) and cocaine (dark blue) sessions. Lighter lines represent averaged velocity across individual days. (**F, G**) Exemplary images of a mouse exposed to saline (left) and cocaine (right) showing differences in pupil dilation. (**H**) Graph representing pupil diameter during cocaine and saline sessions. Graphs represent averages for all mice ± SEM. (**I**) Averaged pupil diameter during saline (light blue) and cocaine (dark blue) sessions. Lighter lines represent the averaged diameter across individual days. (**J, K, M, N**) Exemplary images before and during lip movement (**J, K**) and licking behaviors (**M, N**). (**L, O**) Graphs representing averaged numbers of lip movements (**L**) and licks (**O**) during cocaine and saline sessions. Graphs represent averages for all mice ± SEM. P) Graph representing averaged pupil size 10 seconds before and 20 seconds after a lick of sugar (green) and quinine (yellow). The dashed line represents the moment of a lick. (**R, S**) Graphs showing average pupil size across all trials with sugar (**R**) and quinine (**S**). N numbers in groups: for (**D)** and (**E**): N = 22 and for (**H, I, L, O)**: N = 17, where N = number of animals. Number of groups for **P-S**: N sugar = 20 and N quinine = 15, where N = number of animals. Statistical differences for (**D, H, L, O)** were measured with one-way ANOVA with Dunnett’s multiple comparisons test, with p values for D day 1 vs. day 2 p = 0.0027, day 1 vs. day 3 p < 0.0001, and day 1 vs. day 5 p < 0.0001, and p values for H day 1 vs. day 2 p = 0.0035, day 1 vs. day 3 p = 0.0029, and day 1 vs. day 5 p = 0.0103, and p values for (**L)** day 1 vs. day 3 p = 0.0436, and p values for O day 1 vs. day 3 p = 0.0389. Statistical differences for R and S were measured with the Wilcoxon test, with p values for R p = 0.0385 and for S p = 0.0398. Statistical differences are represented by stars, where *, **, ***, **** correspond respectively to p < 0.05, p < 0.01, p < 0.001 and p < 0.0001.

During imaging, the head-fixed mice were standing on a freely rotating disk that allowed measuring its rotation and indicating movement velocity. With an infrared camera, we recorded mouse facial expressions. During these video recordings, we tracked several points on the mouse face using DeepLabCut software (Fig. 2 C). Finally, we used SimBA software and supervised machine learning (16) to determine several facial movements, such as jaw, nose, or ear movements and licking.

In this highly controlled environment, we identified several behavioral and facial features of the cocaine-intoxicated, head-fixed mice. Firstly, mice exhibited significantly more locomotor activity during cocaine sessions compared to saline sessions, with a peak around 20 minutes after the injection (Fig. 2 D, E). We observed the sensitization pattern (increased behavioral response after repeated exposure to a drug), which is a typical effect of psychostimulants such as cocaine. Another prominent indicator of cocaine’s effect was pupil size, which dilated after cocaine injection, even though mice were in darkness during the procedure (Fig. 2 F-I). We also detected several facial movements and found that cocaine triggered more lip (jaw) movement and licking behavior (Fig. 2 J-O). By calculating relative distances between points, we found other predictors of cocaine’s effects, such as larger angles between mouth, eye, and lip or shorter distances between nose and eye or between nose and whiskers (Fig. Supp. 2 E-J). Other behaviors, such as blinking and whisker movements, were less reliable predictors of cocaine intoxication but mice exhibited more nose movements during cocaine sessions (Fig. Supp. 2 A-D).

To examine mouse facial expressions during drinking sessions, we manually analyzed the video recordings with BehaView software to determine time points when mice were licking droplets. We then analyzed mouse facial expressions 10 seconds before to 20 seconds after the lick. We observed an increased pupil size after both sugar and quinine exposure (Fig. 2 P-S) and a shortened distance between the eye and whiskers directly after licking, which lasted longer after sugar exposure (Fig. Supp. 2 K, L, O, P).

### Opposite activation of DRD1 and DRD2 in the CeM cells after cocaine exposure

We imaged the calcium activity of dopamine-sensitive neurons in the CeM of DRD1-Cre and DRD2-Cre mice through a GRIN lens. For this purpose, we used a custom-made two-photon microscope for calcium imaging of head-fixed mice. In a Cre-dependent manner, we expressed a fluorescently labeled calcium indicator (GCaMP8m) in the CeM of DRD1-Cre or DRD2-Cre mice. We implanted a Gradient Index (GRIN) lens above the CeM, serving as an optical relay lens, allowing imaging neurons located deep inside the brain. For head fixation, a head plate was attached to the mouse skull, ensuring stable immobilization of the animal’s head under the microscope’s objective (Fig. 2 B). Mice were subcutaneously injected with either saline or cocaine, and the calcium activity was imaged for 21 min at 30 Hz, averaged to 10 Hz. In mice with post-mortem validated expression of cre-dependent GCaMP8m and accurate GRIN lens localization, we performed further analysis of calcium events (Fig 3A, D, E).

**Fig. 3:**
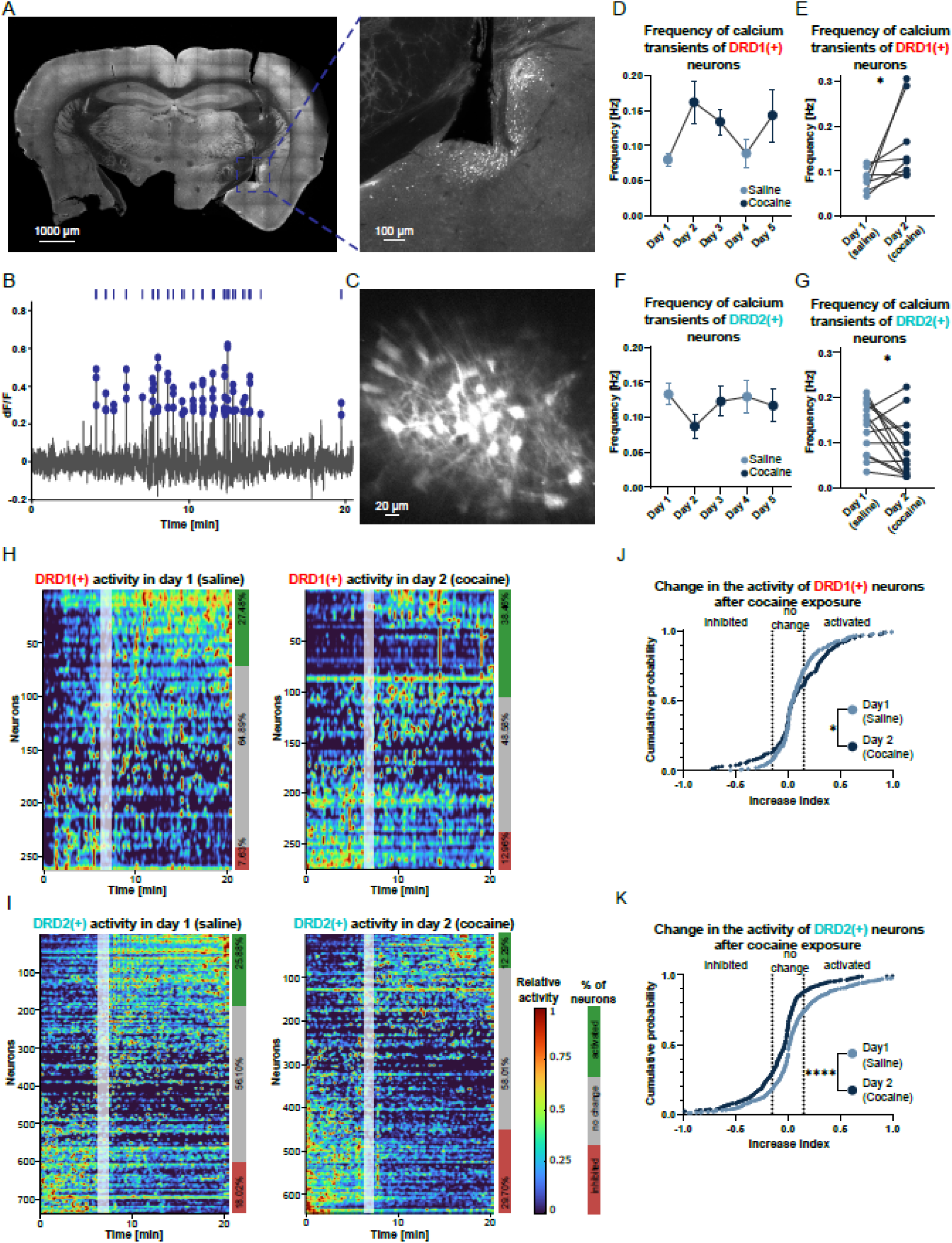
Calcium activity of DRD1- and DRD2-positive neurons in the medial part of the central amygdala during cocaine exposure. (**A**) Example image of the scar in the brain after GRIN lens insertion. Cells expressing GCaMP8m are shown in white. (**B**) Exemplary calcium trace with detected events. (**C**) Example image from a two-photon microscope of GCaMP8m-expressing neurons in the CeM, visible through a GRIN lens. (**D, F**) Graphs showing the frequency of calcium events detected in DRD1(+) and DRD2(+) neurons across all sessions. Graphs represent averages of events for all mice ± SEM. (**E, G**) Graphs showing the averaged frequency of calcium events in DRD1(+) and DRD2(+) neurons during the first saline (day 1) and first cocaine (day 2) injections. Graphs represent individual averages of events for each mouse. (**H, I**) Heatmaps showing calcium activity of all recorded neurons during first saline (day 1, left) and first cocaine (day 2, right) injections for DRD1(+) and DRD2(+) neurons. For each neuron, the activity was binned into 3-second intervals and then normalized to its maximum activity. Neurons were sorted based on the difference in their activity before and after 7.5 minutes of recording (time indicated by a white vertical line). Cells were categorized into “activated” (they increase their activity), “inhibited” (decreased activity), and “no change” (remaining cells). (**J, K**) Graphs representing the distribution of the difference in cell activity before and after 7.5 minutes of recording (Increase index) for DRD1(+) and DRD2(+) neurons. On graphs dotted lines represent the cut-off for cell activity change. For DRD1(+) N = 8 and for DRD2(+) N = 16, where N = number of animals. Statistical differences for (**D**) and (**F**) were measured with one-way ANOVA with Dunnett’s multiple comparisons test. Statistical differences for (**E**) and (**G**) were measured with paired t-tests with p-value p = 0.0407 for DRD1(+) and p = 0.0270 for DRD2(+). Statistical differences for J and K were measured with Kolmogorov-Smirnov test p-value p = 0.0188 for DRD1(+) and p < 0.0001 for DRD2(+). Statistical differences are represented by stars, where *, and **** correspond respectively to p < 0.05 and p < 0.0001.

Our analysis revealed that CeM DRD1(+) and DRD2(+) neurons are opposingly regulated by cocaine (Fig. 3 B, C, F, G). The most striking effects were observed during the first drug exposure, and in DRD1(+) neurons we found an increased frequency of calcium transients, while in DRD2(+) neurons, the frequency decreased. Furthermore, the activity of DRD1(+) cells remained elevated during subsequent cocaine sessions but DRD2(+) cell activity was only affected during the first exposure to cocaine. Then, we plotted the activity of all cells from all mice and we sorted them based on their activity pattern during each session. Cocaine’s pharmacological effects following the subcutaneous injection developed gradually, with a notable breaking point around 7 minutes post-injection, where differences (visible as increased locomotor activity or dilated pupil) between saline and cocaine injections started to be pronounced (Fig. 2 E, I). Thus, we sorted these cells based on their activity before and after the 7-minute time point (Fig. Supp. 3 A, B). Each cell’s activity was normalized to its maximum activity during a session (Fig. 3 H, I). Then, by comparing the activity at the beginning and end of each session, we classified all cells into three categories: activated (relative activity increased by 0.15), inhibited (relative activity decreased by 0.15), and “no change” (remaining cells). The majority of cells fell into the “no change” category and the proportion of activated and inhibited neurons differed for saline and cocaine sessions (Fig. 3 I, K). Moreover, the first cocaine injection caused more DRD1 cells to activate (38.46%) than inhibit (12.96%) while DRD2 cells showed the opposite: 12.29% activated and 29.70% inhibited their activity (Fig. 3 H, I).

### Opposite regulation of DRD1 and DRD2 neurons in the CeM upon sugar and quinine exposure

Even more robust effects were observed when we evaluated the activity of DRD1 and DRD2 cells responding to appetitive and aversive stimuli – sugar and quinine. We measured DRD1 and DRD2 cell activity 10 seconds before and 20 seconds after the lick and calculated the relative activity of each cell based on their maximum activity. On average, DRD1(+) neurons increased and DRD2(+) decreased their activity after sugar consumption (Fig. 4 B, C, E, F). Moreover, DRD2(+) showed a pattern with a peak just before the lick and a dip immediately after. We categorized cells into three classes: activated (relative activity increased by 0.04), inhibited (relative activity decreased by 0.04), and “no change” (remaining cells) (Fig. 4 A, D).

**Fig. 4:**
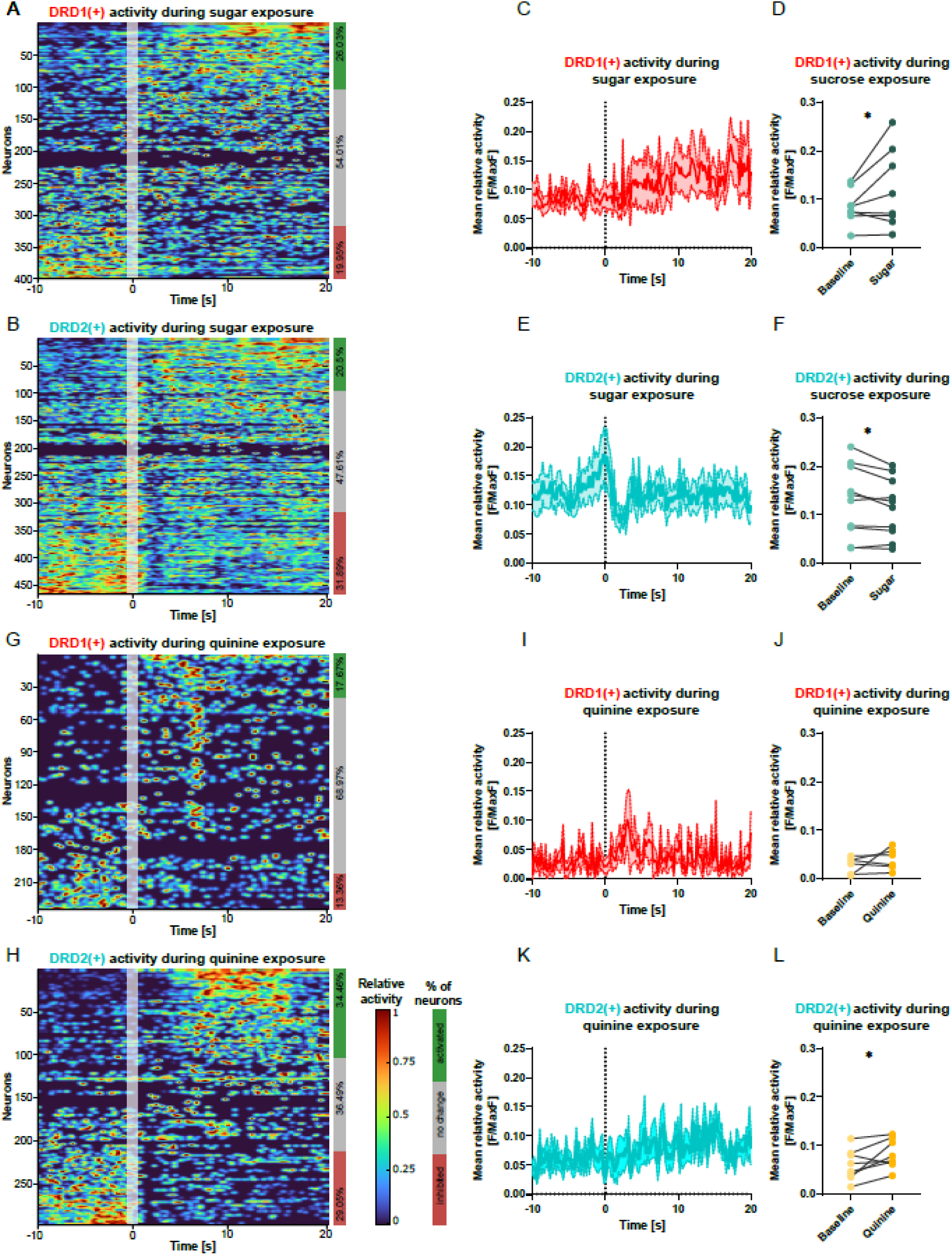
Calcium activity of DRD1- and DRD2-positive neurons in the medial part of the central amygdala during sugar and quinine exposure. (**A, B**) Heatmaps showing calcium activity of all recorded DRD1(+) (**A**) and DRD2(+) (**B**) neurons averaged across all trials for 7.5% sucrose solution exposure. The activity of each neuron was normalized to its maximum activity. Neurons were sorted based on the difference in their activity before and after the lick (the time of the lick is marked with a white line). Cells were categorized into “activated” (they increase their activity), “inhibited” (decreased activity), and “no change” (remaining cells). (**C, E**) Graphs representing relative activity means of DRD1(+) (**C**) and DRD2(+) (**E**) neurons before and after a lick of sucrose solution averaged for all mice. (**D, F**) Graphs showing relative activity means of DRD1(+) (**D**) and DRD2(+) (**F**) neurons before (Baseline) and after a lick of sucrose solution (Sugar). Graphs represent individual averages of the relative activity for each mouse. (**G, H**) Heatmaps showing calcium activity of all recorded DRD1(+) (**G**) and DRD2(+) (**H**) neurons averaged across all trials for 0.01mM quinine solution exposure. The activity of each neuron was normalized to its maximum activity. Neurons were sorted based on the difference in their activity before and after the lick (represented with a white line). Cells were categorized into “activated” (they increase their activity), “inhibited” (decreased activity), and “no change” (remaining cells). (**I, K**) Graphs representing relative activity means of DRD1(+) (I) and DRD2(+) (K) neurons before and after a lick of quinine solution averaged for all mice. (**J, L**) Graphs showing relative activity means of DRD1(+) (**D**) and DRD2(+) (**F**) neurons before (Baseline) and after a lick of quinine solution (Quinine). Graphs represent individual averages of the relative activity for each mouse. N numbers in groups for sugar exposure: DRD1(+) N = 8, for DRD2(+) N = 11. N numbers in groups for quinine exposure: DRD1(+) N = 6, for DRD2(+) N = 8, where N = number of animals. Statistical differences for (**D, F, J,** and **L**) were measured with paired t-tests. For sucrose DRD1(+) Baseline vs. Sugar p = 0.0446, and for sucrose DRD2(+) Baseline vs. Quinine p = 0.0316, and for quinine DRD1(+) Baseline vs. Quinine p = 0.3245, and for quinine DRD2(+) Baseline vs. Quinine p = 0.0471. Statistical differences are represented by stars, where * corresponds to p < 0.05.

Next, we analyzed the activity of neurons during quinine trials. We found that only DRD2(+) neurons increased their relative activity (Fig. 4 G-L). Since mice were exposed to both sugar and quinine in the same imaging session, we could compare the activity of individual neurons during appetitive (sugar) and aversive (quinine) stimuli exposure. We calculated the relative activity of each neuron in both types of trials and found that neurons predominantly responded (with increased or decreased activity) to either sugar or quinine (Fig. 5). Only a small portion of cells responded to both stimuli (Fig. 5 B, D). Moreover, in both DRD1 and DRD2 populations, cells that increased their activity after sugar were inhibited after quinine and vice versa (Fig. 5 A, C).

**Fig. 5:**
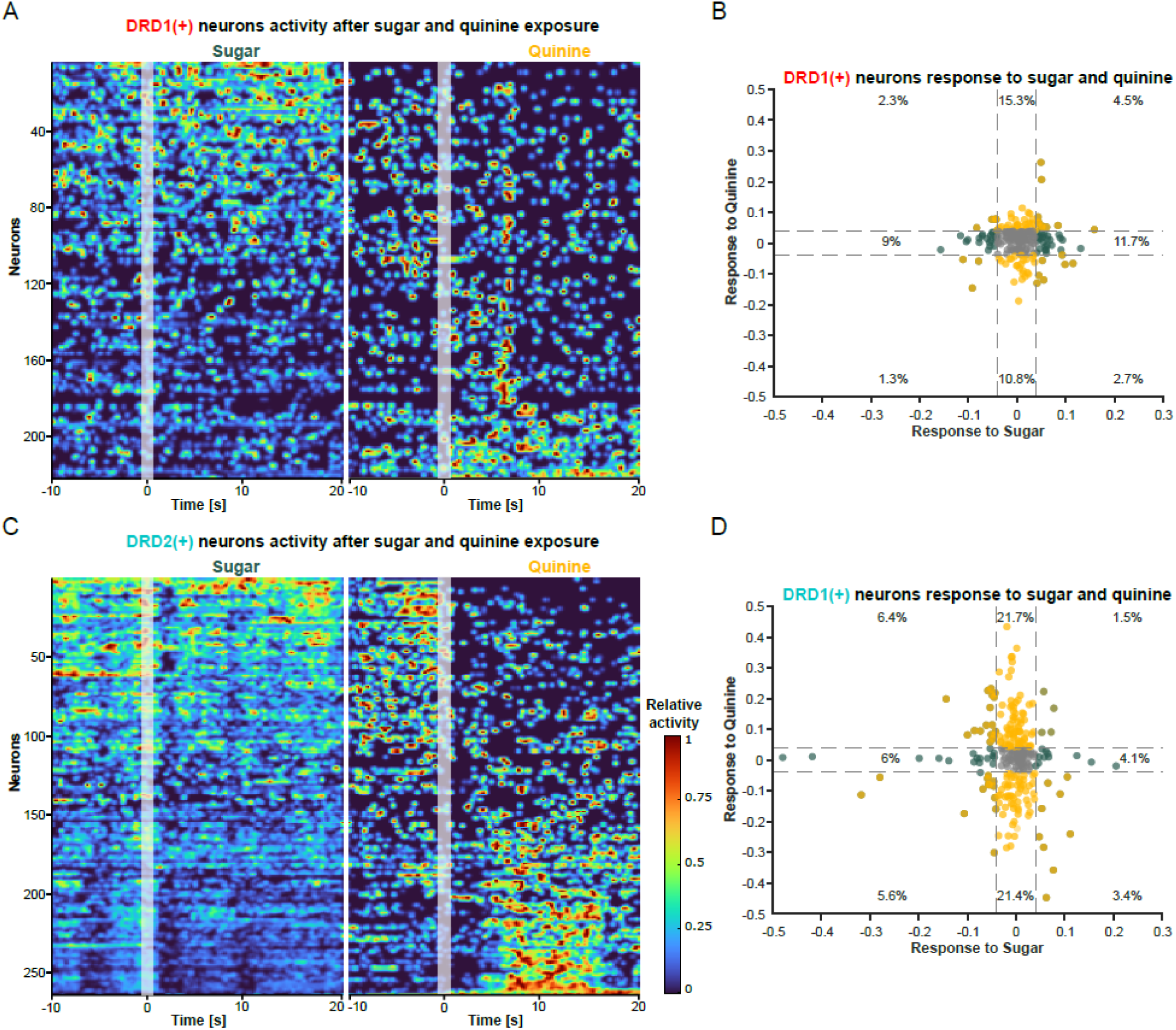
Different populations of DRD1- and DRD2-positive neurons in the medial part of the central amygdala are activated after sugar and quinine exposure. (**A, C**) Heatmaps showing calcium activity of all recorded DRD1(+) (**A**) and DRD2(+) (**C**) neurons. Averaged activity of neurons across all trials for 7.5% sucrose solution and 0.01 mM quinine exposure. The activity of each neuron was normalized to its maximum activity across responses to sucrose and quinine. Neurons were sorted based on the difference in their relative activity after a lick of sucrose solution and a lick of quinine solution. The time of licks is represented as a white line. (**C, E**) Plots representing relative responses of neurons after a lick of sugar (green) and quinine (yellow) for DRD1(+) (**B**) and DRD2(+) (**D**) neurons. Dashed lines represent cut-offs of neuronal responses. Numbers of animals in groups for DRD1(+) N = 6, and for DRD2(+) N = 8.

### Cocaine and sugar induce c-Fos expression in DRD1 and DRD2 neurons in the CeM

Altered activities of dopamine-sensitive neurons in the CeM suggest their involvement in processing information about cocaine and sugar. We wanted to investigate whether this altered activity triggers lasting plasticity processes. Therefore, we examined the expression of synaptic plasticity markers in DRD1(+) and DRD2(+) neurons in the CeM of mice exposed to cocaine or sugar. We employed a seven-day reward exposure protocol, previously shown to induce plastic changes in the NAC and the CeA ^6,10^.

In this study, DRD1-TdTomato/DRD2-GFP mice received intraperitoneal (i.p.) injections of cocaine for seven consecutive days (Fig. 6 A). During this period, the locomotor activity of mice was recorded for 30 minutes using a camera positioned above the cage. The recorded videos were later analyzed with DeepLabCut software to calculate the locomotor activity of mice. Analysis revealed that mice exposed to cocaine exhibited increased locomotor activity and demonstrated sensitization (Fig. 6 C). Compared to mice on a rotating disc from a two-photon experiment, the velocity pattern increased more rapidly, with a peak effect occurring already 1.5 minutes after the injection, demonstrating that subcutaneous and i.p. injections result in different dynamics of cocaine’s effects (Fig. 6 D).

**Fig. 6:**
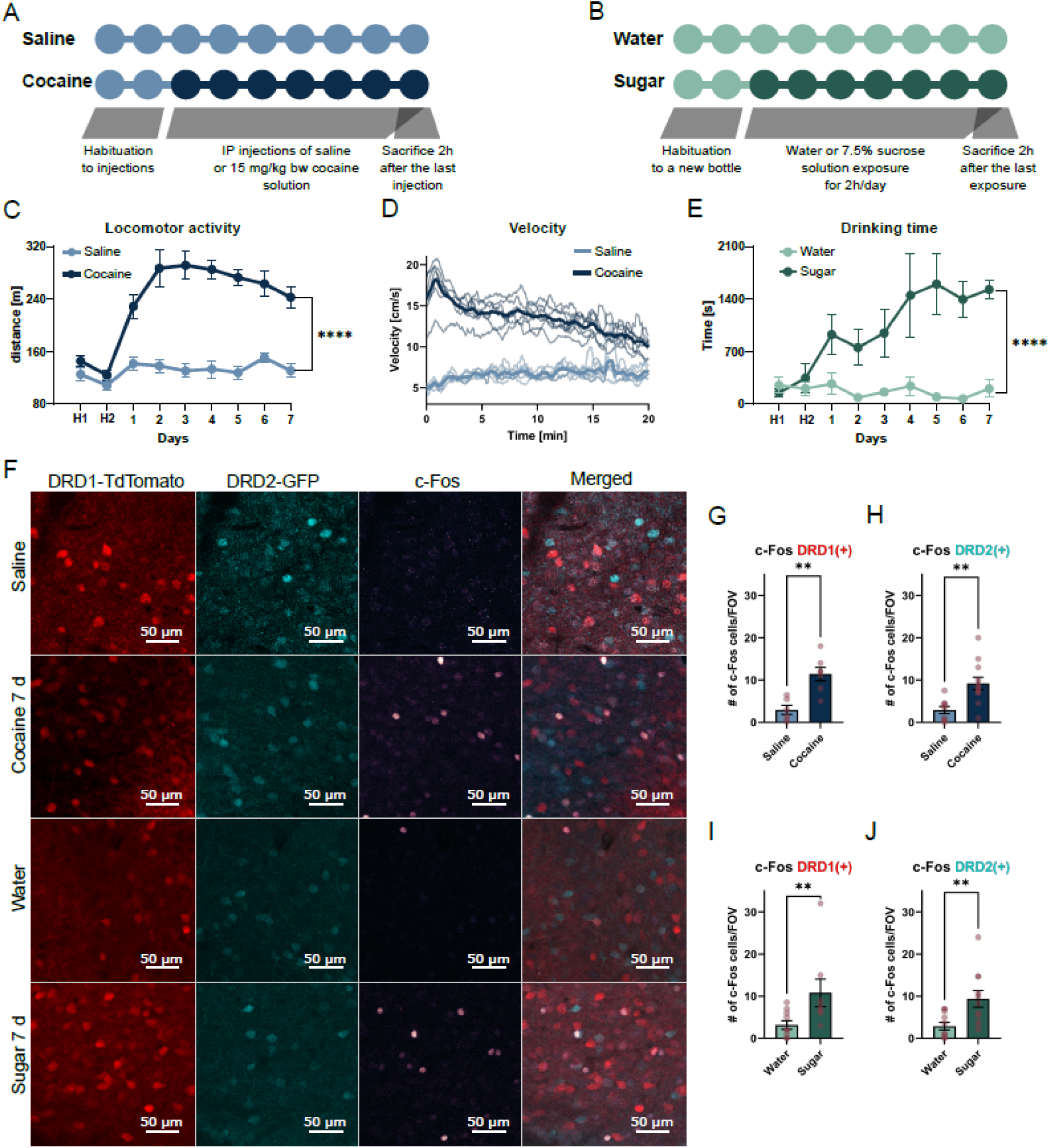
c-Fos expression in the central amygdala of freely-moving mice exposed to cocaine and sugar. (**A, B**) Experiment design. For cocaine exposure (**A**) mice received once daily intraperitoneal injections of saline or 20 mg/kg body weight cocaine solution. For sugar exposure (**B**) a bottle with fresh water was replaced with 7.5% sucrose solution (or another fresh water bottle for the control group) for 2 hours per day. Two hours after the last reward exposure mice were sacrificed. (**C**) Graph representing locomotor activity of a mouse in a cage for 30 minutes after cocaine and saline injections. The graph represents averages for all mice ± SEM. D) Averaged velocity of mice running in a cage for 30 minutes after cocaine (dark blue) and saline (light blue) injections. Lighter lines represent averaged velocity across individual days. (**E**) Graph showing time spent on drinking water or 7.5% sucrose solution (Sugar) during 2-hour exposure sessions. (**F**) Exemplary images of c-Fos-expressing neurons in the central nucleus of the amygdala of DRD1-TdTomato/DRD2-GFP mice exposed to saline, cocaine, water, and sugar. (**G, H, I, J**) Graphs representing an average number of c-Fos-positive cells in CeM in DRD1(+) and DRD2(+) neurons after cocaine (**G, H**) and sugar exposures (**I, J**). Graphs represent averages ± SEM. Red dots represent values for individual mice. N number in groups for (**C, D, E**): N Saline = 7; N Cocaine = 7, N Water = 6; N Sugar = 7 where N = number of animals. N number in groups for (**G-J**): DRD1(+) N Saline = 6(18); DRD1 (+) N Cocaine = 7(8), DRD2(+) N Saline = 9(22); DRD2(+) N Cocaine = 12(20), DRD1(+) N Water = 10(14); DRD1(+) N Sugar = 8(13) and for DRD2(+) N Water = 11(19); DRD2(+) N Sugar = 11(25), where N = number of animals (number of analyzed ROIs). Statistical differences for (**C**) and (**E**) were determined with Two-way ANOVA with repeated measurements, where p-value < 0.0001. Statistical differences for (**G**) and (**H**) were measured with unpaired t-tests, where for DRD1(+) Saline vs. Cocaine p-value = 0.0012 and for DRD2(+) Saline vs. Cocaine p-value = 0.0025. Statistical differences for (**I**) and (**J**) were determined with the Mann-Whitney Test, where for DRD1(+) Water vs. Sugar p-value = 0.0077 and for DRD2(+) Water vs. Sugar p-value = 0.0062. Statistical differences are represented with stars, where ** and **** correspond respectively to p < 0.01 and p < 0.0001.

To evaluate the effects of sugar, another set of mice received access to a bottle containing sweet water (7.5% sucrose solution) for 2 hours per day over seven consecutive days (Fig. 6 B). This bottle was connected to a custom-made lickometer to measure time spent on drinking. Similar to previous observations, we noticed a strong preference for sweet water (Fig. 2 E and Fig. 6 E) ^6,10^.

Two hours after the onset of the final reward exposure, mice were sacrificed and perfused with 4% paraformaldehyde in PBS. Their brains were sliced on a vibratome and slices were labeled for c-Fos protein, a marker of neuronal activity related to synaptic plasticity ^22–24^. Using a confocal microscope, we imaged the CeM and observed that both rewards increased c-Fos levels in both DRD1(+) and DRD2(+) neurons (Fig. 6 F-J). Although most imaged slices came from single transgenic mice labeled for either DRD1(+) or DRD2(+) neurons, some slices from double-transgenic animals allowed us to compare c-Fos levels in dopamine-sensitive cells to the entire population of c-Fos-positive neurons in the CeM. Despite the limited number of double-labeled samples, our analysis indicated that dopamine-sensitive cells constituted less than 50% of all c-Fos-positive cells, indicating that other, neuronal populations in CeM were also activated during sugar and cocaine exposures (Fig. Supp. 4 B, C).

### Cocaine and sugar exposure induce opposing plastic changes in DRD1(+) and DRD2(+) neurons in the CeM

Immediate early gene expression products are useful tools for identifying cells undergoing plasticity changes in response to various stimuli. However, the manner of plasticity changes can involve the strengthening or weakening of synaptic connections between neurons, which can be studied by examining the electrical properties of neurons. Therefore, in another set of experiments, we performed ex vivo whole-cell patch clamp experiments on brain slices from DRD1-TdTomato/DRD2-GFP mice.

For that, two hours after the final exposure to cocaine or sucrose, mice were sacrificed, and their brains were sliced using a vibratome. DRD1(+) and DRD2(+) neurons were identified based on their endogenous fluorescence, and spontaneous Excitatory Postsynaptic Currents (sEPSCs) were recorded to assess their excitatory (AMPA receptors-mediated) drive on recorded neurons (Fig. 7 A). We found that cocaine opposingly altered the sEPSCs frequency of DRD1(+) and DRD2(+) neurons: cocaine-treated mice had increased sEPSCs frequency in DRD1(+) and decreased in DRD2(+) neurons (Fig. 7 B, D-G). However, we did not observe any changes in sEPSCs amplitude (Fig. 7 H-K). Interestingly, we found similar opposing alterations in the sucrose-treated group. In this case, however, DRD2(+) neurons showed increased sEPSCs frequency, while DRD1(+) neurons showed a decreased frequency (Fig. 7 C, L-O). Again, the amplitude of these neurons was not affected by the reward (Fig. 7 P-T).

**Fig. 7:**
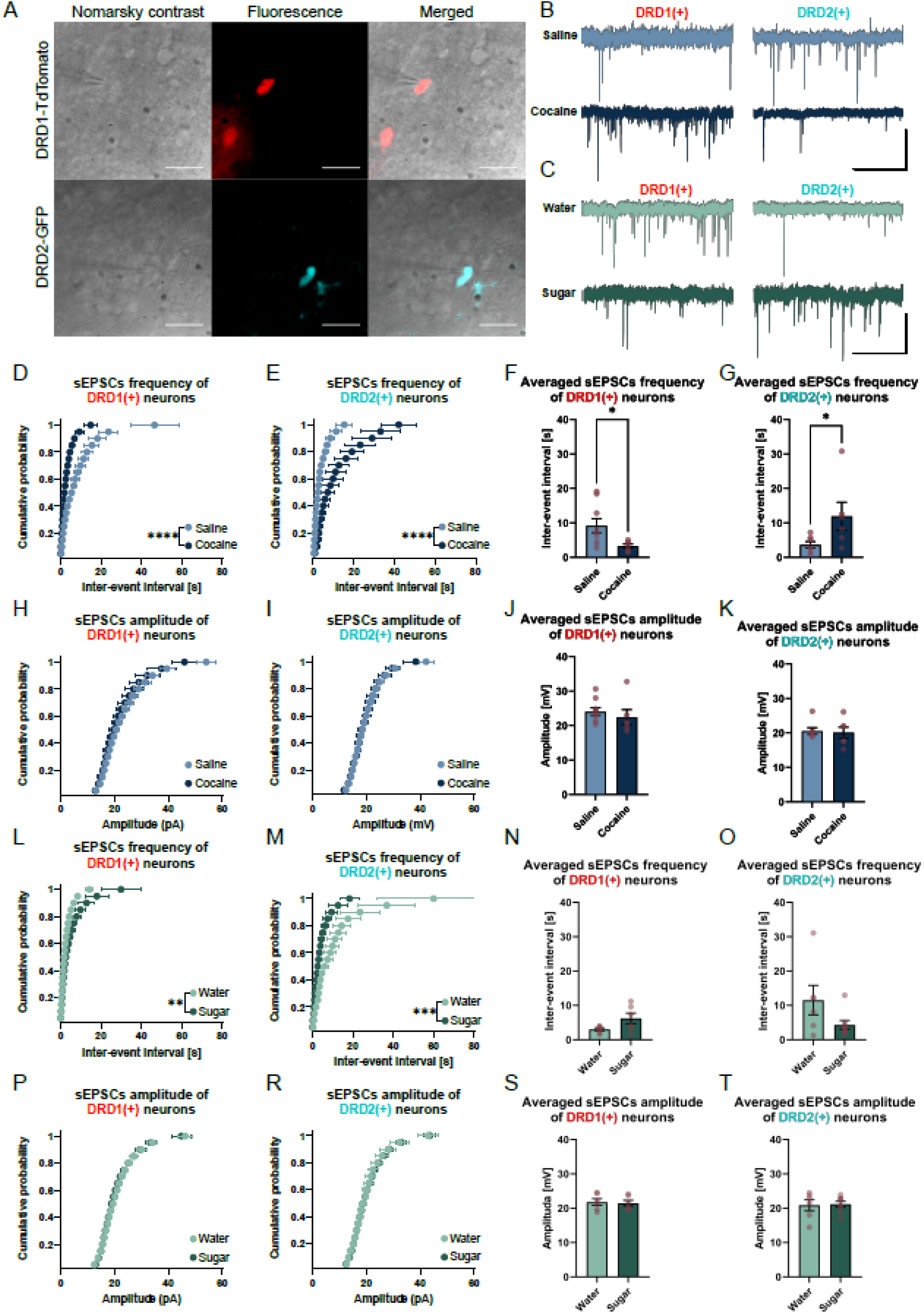
Electrophysiology recordings from DRD1- and DRD2-positive neurons from the medial nucleus of the central amygdala in mice exposed to cocaine or sugar. (**A**) Exemplary images of DRD1-TdTomato (red) and DRD2-GFP (cyan) neurons; scale bar 20 μm. (**B, C**) Example recordings of sEPSCs from DRD1(+) and DRD2(+) neurons of mice exposed to saline/cocaine (blue, **B**) or water/sugar (green, **C**). Horizontal bars represent 2 s and vertical 20 pA. (**D, E, L, M**) Graphs show the cumulative distribution of inter-event intervals of DRD1(+) and DRD2(+) neurons after cocaine (**D, E**) or sugar exposure (**L, M**). Graphs represent averages ± SEM. (**F, G, N, O**) Graphs show averaged sEPSCs inter-event intervals of DRD1(+) and DRD2(+) neurons after cocaine (**F, G**) or sugar exposure (**N, O**). Graphs represent averages ± SEM. Red dots represent values for individual mice. (**H, I, P, R**) Graphs show the cumulative distribution of amplitudes of events of DRD1(+) and DRD2(+) neurons after cocaine (**H, I**) or sugar exposure (**P, R**). Graphs represent averages ± SEM. (**J, K, S, T**) Graphs show averaged sEPSCs amplitude of DRD1(+) and DRD2(+) neurons after cocaine (**J, K**) or sugar exposure (**S, T**). Graphs represent averages ± SEM. Red dots represent values for individual mice. N number in groups for cocaine exposure: N DRD1(+) Saline = 9(29), N DRD1(+) Cocaine = 6(15), N DRD2(+) Saline = 7(17), N DRD2(+) Cocaine = 6(24). N number in groups for sugar exposure: N DRD1(+) Water = 6(30); N DRD1(+) Sugar = 6(17), N DRD2(+) Water = 6(33); N DRD2(+) Sugar = 8(29), where N = number of animals (number of recorded cells). Statistical differences for (**D, E, H, I, L, M, P)**, and (**R**) were measured with the Kolmogorov-Smirnov test, where for (**D**) and (**E**) p-value < 0.0001, for (**L**) p-value = 0.0029, and for M p value = 0.0003. Statistical differences for (**F**) were measured with an unpaired t-test, where p-value = 0.0419. Statistical differences for (**G, J, K**), and (**N-T**) were measured with the Mann-Whitney Test, with p-value = 0.0350 for (**G**). Statistical differences are represented by stars, where *, **, ***, **** correspond respectively to p < 0.05, p < 0.01, p < 0.001 and p < 0.0001.

During electrophysiology experiments, biocytin was added to an internal solution used for patch clamping. This approach allowed us to later label cells with streptavidin conjugated with fluorescent dye, to visualize the dendritic tree, and to calculate dendritic spine density (Fig. Supp 5 A, D). We found no changes in spine density neither in DRD1(+) nor DRD2(+) cells, after either of the rewards exposures (Fig. Supp. 5 B, C, E, F).

### Modulation of neither DRD1 nor DRD2 neurons in the CeM affects cocaine-conditioned place preference

To investigate whether modulating dopamine-sensitive neurons in the CeM will affect cocaine-associated memories, we chemogenetically manipulated the activity of these neurons during the conditioned place preference (CPP) test. DRD1-Cre and DRD2-Cre mice expressed the cre-dependent DREADD construct (excitatory hM3Dq, inhibitory hM4Di, or sham for control mice) in the CeM (Fig. 8 A). The mice then underwent the CPP procedure, during which they developed a preference for the cocaine-associated chamber (Fig. 8 A, D). After the conditioning phase, on the 10th day, DREADD receptors were activated via i.p. injections of CNO. All cocaine-treated groups showed a preference for cocaine-associated chambers. However, neither activation nor inhibition of dopamine-sensitive neurons affected the mice’s preference. Thus, manipulation of either DRD1(+) or DRD2(+) cells in the CeM does not affect the recall of cocaine-associated memories or the preference execution (Fig. 8 B, C).

**Fig. 8:**
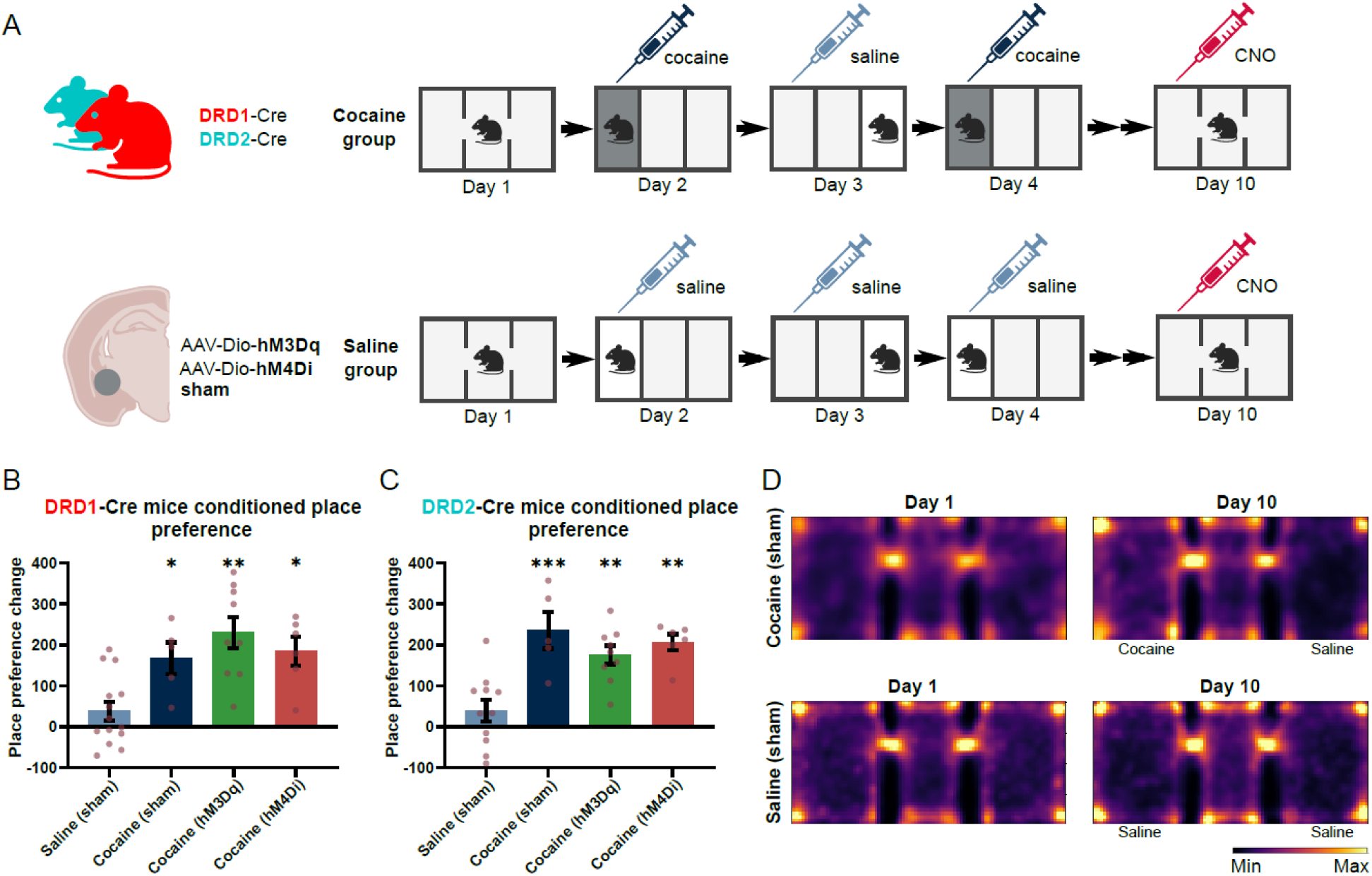
Cocaine-induced conditioned place preference of mice with manipulated activity of dopamine-sensitive neurons in the CeM. (**A**) Scheme of the experiment. Excitatory DREADD (hM3Dq) or inhibitory DREADD (hM4Di) were expressed in DRD1-Cre and DRD2-Cre mice. After the recovery, mice were introduced to a three-chamber cage. In the next 9 days, they received either saline or cocaine i.p. injection. During the 10th day, mice received CNO i.p. injection 30 minutes before the test. (**B, C**) Place preference change of DRD1-Cre (**B**) and DRD2-Cre (**C**) mice. (**D**). Heatmaps were created based on the average time spent in the arena of all mice from the control group (without DREADD expression). All traces were transformed in the way that the left chamber is a chamber where mice received cocaine i.p. injections. Number of groups for DRD1-Cre mice: N Saline (sham) = 14, N Cocaine (sham) = 5, N Cocaine (hM3Dq) = 9, N Cocaine (hM4Di) = 6. Number of groups for DRD2-Cre mice: N Saline (sham) = 11, N Cocaine (sham) = 5, N Cocaine (hM3Dq) = 9, N Cocaine (hM4Di) = 6. Statistical differences were measured with Two-way ANOVA with repeated measurements, where for DRD1-Cre Saline (sham) vs. Cocaine (sham) p-value = 0.05, Saline (sham) vs. Cocaine (hM3Dq) p-value =0.002, Saline sham) vs. Cocaine (hM4Di) p value = 0.0441 and for DRD2-Cre Saline (sham) vs. Cocaine (sham) p value = 0.001, Saline (sham) vs. Cocaine (hM3Dq) p value = 0.0042, Saline (sham) vs. Cocaine (hM4Di) p value = 0.032. Statistical differences are represented by stars, where *, **, and ***, correspond respectively to p < 0.05, p < 0.01, and p < 0.001.

## Discussion

The CeA is an immensely complex structure with diverse inputs and cell types, reflecting its key function in processing and interpreting emotional responses in various behaviors. Growing evidence supports its role in the development of addiction to substances, such as alcohol, opioids, or stimulants ^25,26^. Although the CeA is not traditionally considered a primary site of dopaminergic activity, recent research has shown that dopamine signaling plays a crucial role in modulating emotional responses, stress, and addiction-related behaviors.

### DRD1 and DRD2 cells distribution in the CeA

Studies in the ’90s described the central amygdala of the rat as being populated mainly by the DRD2-expressing cells, while the BLA was predominantly expressing DRD1 ^27,28^. More recently, in RNA sequencing studies in mice, DRD1(+) cells were assigned to intercalated cells of the amygdala, CeM, and CeL nuclei, while DRD2(+) cells were found in the CeC, CeM, and CeL ^29–31^. On the whole, our data indeed suggests that the CeA is mainly populated with DRD2(+) cells, with the CeL and CeC nuclei almost completely devoid of DRD1(+) neurons. However, in the CeM nucleus, both types of neurons are present, in similar numbers, and amount to nearly 40% of all cells there.

The RNA sequencing studies revealed that the CeA is a brain region rich with diverse cell types, each characterized by specific receptors or proteins that define their physiological role. In this study, we colocalized dopamine-sensitive cells with PKCδ, a protein especially abundant in CeL neurons ^32^. PKCδ(+) neurons in the CeL are known to participate in defensive behaviors, while cocaine i.p. injections, as well as DRD2 agonist, increase the number of c-Fos(+) in the CeL ^33^. Similarly to the earlier report ^32^, we showed only a partial overlap of DRD2 and PKCδ, with the majority of cells expressing one marker or the other. Our in vivo imaging of rewards processing shows that cocaine and sugar, though in general predominantly increase the activity of DRD1(+) cells and decrease DRD2(+), engage both populations. Moreover, the percentage of dopamine-sensitive neurons among all c-Fos(+) cells was less than 50%. It indicates that dopamine-sensitive cells were not the only subpopulation participating in cocaine or sugar processing in the CeM. Thus, the genetic profile of a neuron does not yet determine its role in how the amygdala processes information.

Investigating structures that send their projections to dopamine-sensitive cells in the central amygdala, we discovered DRD1(+) cells as the main input source. We observed that CeM receives DRD1(+) cell inputs from the sensory and prefrontal areas of the cortex (PFC). Cortical DRD1-expressing cells were mainly studied in the context of PFC-BLA projection that participates in decision-making ^34^ and the retrieval of extinguished fear ^35^. One study shows that the infralimbic (IL) PFC DRD1(+) to CeM pathway mediates the reinstatement of fear. Blocking D1 receptors in the IL prevented c-Fos expression in the CeM and reduced freezing behavior triggered by the reminder foot shock ^36^. D1-expressing interneurons in the medial PFC provide inhibition to the BLA and CeA. Dopamine reduces their excitability which results in shifting the balance from inhibition to excitation in the BLA and CeA ^37^.

Interestingly, we found strong retrograde labeling in the ZI. Recent reports identified the ZI as a structure strongly implicated in fear memory and nociception ^38–41^. Dopaminergic neurons that reside there and project to the BLA were shown to affect aversive learning in a social defeat stress model ^40^. Inhibitory projections from the CeA to the ZI participate in fear conditioning and recall ^41^, while chemogenetic manipulations of PKCδ(+) neurons projecting from the CeL and CeC to ZI changed sensitivity to tactile and painful stimuli ^42^.

### DRD1 and DRD2 cells function in reward processing in the CeA

The role of D1- and D2-expressing neurons in the amygdala has been mainly studied by injecting dopamine receptor agonists/antagonists via implanted cannulas. Those approaches, therefore, affect the entire amygdala complex, not its individual subdivisions. In the aspect of feeding behavior, it was reported that activation of D2-expressing cells reduced food intake and sucrose operant conditioning in rats, while inhibition or activation of D1-expressing cells didn’t have much effect ^43^. On the other hand, D1-receptor blocking in the CeA reduced orexin-strengthened saccharin preference ^44^. In other studies, activation of D2 cells in the central amygdala showed their involvement in rewarding pain relief properties. As the level of the D2 receptor is decreased, activating D2 cells with an agonist reverses pain relief ^45^. Others showed the involvement of dopamine-sensitive cells in cocaine withdrawal, which was shown to increase the D1 receptor level in the CeL after 2 weeks of abstinence from cocaine ^46^, while activation of D2 receptors (but not D1) decreased cocaine self-administration and cocaine-induced reinstatement of seeking behavior ^47^.

Focusing specifically and separately on D1- or D2-type cell populations, we show that in the CeM, DRD1(+) cells differ from DRD2(+) in their synaptic response to sugar and cocaine. With no effect on spine density in DRD1- and DRD2-expressing populations, the observed change in the frequency of spontaneous EPSCs points to a presynaptic mechanism such as changes in the probability of neurotransmitter release. Our data suggests that extended (7-day) exposure to i.p. cocaine injections increases glutamatergic drive to DRD1(+) neurons and decreases to DRD2(+). Sugar exerts the exact opposite effect – after 7-day exposure, DRD2(+) neurons would receive stronger excitatory input than DRD1(+) cells. Overall, without distinguishing different neuronal populations, our previous study reported an increase in the number of silent synapses – immature glutamatergic connections that were triggered by both sugar and cocaine exposures ^10^. Their appearance and subsequent maturation would increase glutamatergic transmission to the CeM neurons. Here we report that presynaptic adaptations depend on the cell type, and rewards shift the balance of how DRD1(+) and DRD2(+) cells would be engaged in their processing.

An increase of sEPSCs frequency in DRD2(+) neurons (suggestive of increased release probability) after repeated sugar exposure is intriguing when paired with the *in vivo* imaging results of mice drinking sucrose and quinine for the first time. Our imaging result showed that sugar decreased the activity of DRD2(+) cells and the majority of DRD2(+) cells reacted to quinine rather than sugar. Perhaps after repeated exposure to sugar, the increase in sEPSCs frequency in DRD2(+) cells is more of a compensation mechanism for their initially reduced activity.

Our chemogenetic manipulation of dopamine-sensitive neurons in the CeM during the expression of CPP (on the final day of the protocol, when the mouse freely explores both chambers) did not alter mice’s preference for the cocaine-paired compartment. Given how dopamine function is crucial in processing novel stimuli, perhaps CNO administration during the conditioning phase, when preference is established, would show differences in DRD1(+) or DRD2(+) cells engagement. However, rats receiving intraamygdalar injections of D1 or D2 receptor antagonists during the conditioning phase did not show changes in their cocaine-induced CPP. An increase in preference was observed when animals were pretreated with neuromodulator neurotensin – an effect that was blocked by D1 and D2 receptor antagonists ^48^. A more possible explanation for the lack of effects of chemogenetic activation/inhibition of DRD1 and DRD2 cells on cocaine-conditioned preference is the fact that (based on our *in vivo* observations and c-Fos expression) only a fraction of all dopamine-sensitive neurons change their activity patterns after cocaine exposure, while plasticity markers are induced in other neuronal populations as well. Identifying and targeting a smaller fraction of both D1 and D2-expressing neurons that belong to one functional circuit would be a better strategy.

### Facial expressions analysis

We performed a comprehensive analysis of mice’s facial expressions in response to sweet/bitter water or cocaine exposure. Similar analyses have been previously done for other stimuli, including bitter and sweet water exposure, painful electric shocks, or unexpected sounds, and were successfully used as a readout of the internal states of animals ^49,50^. For instance, exposure to sugar and quinine, elicited similar facial expressions in mice, primarily reflected in ear positioning and to a lesser extent as a grimace in the lower frontal part of the head ^49^. Unfortunately, in our studies, ear movements could not be tracked, as they were covered by the mounting bar for head fixation. Nevertheless, we found that monitoring the frontal part of the mouse head was sufficient to detect changes in the mouse’s facial expression after exposure to sweet or bitter tastes. Moreover, our study revealed that both sucrose and quinine induced pupil dilation, which is a response commonly recognized as a metric of animal arousal and uncertainty ^50,51^. In our research, pupil dilation was also the most pronounced indicator of cocaine intoxication, an effect known to be triggered by stimulants in humans and used as a rapid screening tool for cocaine use ^52^. Pupil dilation was also observed in rodents before but in our study, we showed that this effect can be observed even when animals’ eyes had adapted to complete darkness ^53^. In our study, we also used tracking points on the mouse head to detect specific movements of the animal under cocaine intoxication. Among the analyzed behaviors, particularly interesting were increased licking and jaw movements, which are also characteristic effects of a psychostimulant overdose in humans, colloquially known as gurning.

Facial expressions and movements in mice may serve as reliable indicators of rewarding stimuli. However, it is uncertain whether the alterations observed in response to cocaine reflect pleasant sensations associated with drug effects, or if they are nonspecific manifestations of impaired circuitry controlling muscle movements. Since the responses to cocaine are similar in both humans and rodents, it raises the possibility that similar pathways are affected. Especially, as both species have a similar organization of facial muscles, which receive inputs from a motor control network in the brainstem, bypassing the spinal cord ^54^.

Facial movements might thus be a valuable indicator of drug-induced behavioral changes, which so far, have been limited to easily observed responses, like tail raising, increased locomotor activity, or fur ruffling. However, whether these facial expressions can reliably serve as a measure of cocaine intoxication remains to be determined, especially by studying whether such effects scale with dosage.

Creating a library will require a more standardized approach to recording and analyzing facial expressions. Current analysis methodologies vary widely, utilizing techniques such as counting occurrences of gradient orientation in localized portions of an image (histogram of oriented gradients, HOG) or tracking individual points on the mouse head ^49,55–57^. Recording techniques also differ, ranging from the most popular side view, through the top view to more sophisticated methods employing multiple cameras for 3D reconstruction of the mouse head ^54,56^.

## Acknowledgments

We thank Prof. Benjamin Judkewitz (Einstein Center for Neurosciences, Charité Universitätsmedizin Berlin, Berlin, Germany) for the help in building the custom-made two-photon microscope. We thank Renata Zakrzewska and Tomasz Włodarczyk (Laboratory of Behavioral Models, Nencki Institute of Experimental Biology PAS, Warsaw, Poland) for their help in the CPP experiment. We thank Urszula Szachowicz (Laboratory of Neuronal Plasticity, Nencki Institute of Experimental Biology PAS, Warsaw, Poland) for her excellent technical support.

Confocal imaging was performed at the Laboratory of Imaging Tissue Structure and Function, which serves as an imaging core facility at the Nencki Institute of Experimental Biology and is part of the infrastructure of the Polish Euro-BioImaging Node. The Polish Euro-BioImaging Advanced Light Microscopy Node is financed by the Minister of Education and Science based on contract no. 2022/WK/05.

## Funding

This study was supported by: National Science Centre, Poland, grant number: 2020/37/N/NZ4/02888 (ŁB) National Science Centre, Poland, grant number: 2023/50/E/NZ4/00421 (AB)

## Author contributions

Conceptualization: ŁB, AB Methodology: ŁB, PS, MP, RŁ, AB Investigation: ŁB, PS, JW, MP, KH Visualization: ŁB, KH, JW Supervision: AB, RŁ Writing—original draft: ŁB, AB Writing—review & editing: ŁB, AB, JW, KH

## Competing interests

Authors declare that they have no competing interests.

## Materials and Methods

### Animals

All experiments were performed on adult, DRD1-TdTomato/DRD2-GFP (B6.Cg-Tg(Drd1a- tdTomato)6Calak/J, RRID: IMSR_JAX:016204, crossed with B6.FVB(Cg)-Tg(Drd2- EGFP)S118Gsat/KreMmucd, RRID: MMRRC_036931-UCD), DRD1-Cre (B6;129-Tg(Drd1- cre)120Mxu/Mmjax, RRID: MMRRC_037156-JAX), or DRD2-Cre (B6.FVB(Cg)-Tg(Drd2-cre)ER44Gsat/Mmucd, RRID: MMRRC_032108-UCD) mice of both sexes, 3–5 months old. Mice were housed in individual cages under a 12 h/12 h light/dark cycle with food and water ad libitum. All studies were performed following the European Council Directive of November 24, 1986 (86/609/EEC), Animal Protection Act of Poland, and approved by the 1st Local Ethics Committee in Warsaw (permission number 463/2018 and 1225/2021). All efforts were made to minimize the number of animals and their suffering.

### Immunolabeling and confocal imaging

Mice were perfused with cold PBS followed by 4% PFA dissolved in PBS. Their brains were removed and kept in 4% PFA for 24 hours in the 4°C and later in 4°C PBS. Brains were sliced on a vibratome (Leica VT 1200 S) into 100 μm slices. Brain slices from DRD1-TdTomato/DRD2- GFP mice were permeabilized at room temperature in 0.25% Triton X-100 in PBS and then incubated for three hours in a blocking solution (3% BSA and 0.1% Triton X-100 in PBS). Slices were then incubated for 24 hours at 0°C in 0.1% Triton X-100 and 0.1% BSA in PBS containing a mix of antibodies against endogenous fluorophores: anti-GFP (Abcam ab13970, 1:1500) and anti-RFP (ThermoFisher M11217, 1:1500). Additionally, a mix contained antibodies against either c-Fos (Synaptic Systems 226 003, 1:2000) or PKCδ (Abcam ab182126, 1:1500). Later, brain slices were incubated for three hours at room temperature with a mix of secondary antibodies (ThermoFisher A-11039 and A32733, 1:400 and Abcam ab150158, 1:400) that had been previously centrifuged at 20000g for 10 minutes. Brain slices from DRD1-Cre and DRD2- Cre for Cre localization were incubated only with anti-Cre antibody (1:3000 ^58^, produced in Division of Molecular Genetics DKFZ) and with secondary antibody ThermoFisher A32733 that had been previously centrifuged at 20000g for 10 minutes.

Brain slices were placed on coverslips with a mounting medium (Fluoromont-G-DAPI) and imaged using Zeiss LSM 780 confocal microscope with 10x (dry, NA = 0.3) and 40X (oil, NA = 1.4) objectives. Diode lasers (405 nm and 561 nm) and an argon laser (488 nm) were used for fluorescence excitation. Signals were collected at the spectra 340-350 nm, 510-530 nm, and 575- 590 nm. Images were later analyzed with a Cell-Counter plugin in FIJI software.

### Electrophysiology and neuronal dendritic spine density measurements

Two hours after the last reward exposure, mice were sacrificed by decapitation. Their brains were removed and sliced in 0°C NMDG solution (135 mM NMDG, 1.2 mM KH2PO4, 1 mM KCl, 1.5 mM MgCl2, 0.5 mM CaCl2, 10 mM D-glucose, 20 mM choline bicarbonate) saturated with carbogen (5% CO2 and 95% O2) using a vibratome (Leica VT 1200 S). 250 μm brain slices were incubated in 31°C ACSF solution (119 mM NaCl, 2.5 mM KCl, 26 mM NaHCO3, 1.3 mM MgCl2, 1 mM NaH2PO4, 20 mM D-glucose; 2.5 mM CaCl2) for 10 minutes, and then for at least another hour in room temperature ACSF solution saturated with carbogen.

Electrophysiology experiments were performed in a carbogen-saturated ACSF solution with 100 μM picrotoxin to block inhibitory transmission. Borosilicate electrodes (4-6 MΩ) were filled with an internal solution (pH 7.1, osmolarity 290-295 mOsm, 120 mM K-gluconate, 2 mM MgCl2, 58 0.4 mM EGTA, 0.1 mM CaCl2, 10 mM HEPES, 2.5 mM Na2-ATP, 0.25 mM Na3-GTP) containing also 5 mM biocytin. All recordings were performed using Igor PRO (Wavemetrics) and a setup equipped with a universal amplifier (npi ELC03XS) and Multi-Channel Data Acquisition Interface (ITC-18, InstruTECH/HEKA). Signals were recorded at 10 kHz and filtered at 2 kHz.

In the voltage-clamp mode, the membrane potential was set to -60 mV and sEPSCs were recorded for 20 minutes to be later analyzed using Clampfit (10.3) software. After each sEPSC recording, in current-clamp mode, the cell was stimulated for 10 minutes with 20 pA current to spread the biocytin within the cell. Brain slices were then fixed for 30 minutes in 4% PFA in PBS at 4 and labeled with streptavidin conjugated with Alexa Fluor 568 (ThermoFisher S11226) for three hours at room temperature. Labeled brain slices were placed on coverslips and imaged using a Zeiss Laser Scanning Microscope 780 with a 63x objective (561 nm argon laser and 575- 590 nm filters). Images were taken at 1024 x 1024 px with a 1.15 μm z-step and analyzed using a Dendritic Spine Counter, a plugin for Fiji software.

### Viral injections and GRIN lens implantation for calcium imaging

DRD2-Cre or DRD1-Cre mice were anesthetized with isoflurane and received a cocktail of painkillers (butorphanol 3 mg/kg, and tolfenamic acid 3 mg/kg). During anesthesia, the skin above the scalp was removed, and the skull was scratched with a scalpel. 200 nl of adeno-associated viruses (pGP-AAV-syn-FLEX-jGCaMP8m-WPRE ^59^ 7×10¹² vg/mL, Addgene 162378-AAV1) were stereotactically injected into the CeM (coordinates from bregma: ML +2.2, AP -1.3, DV 4.6 mm) at a rate of 75 nl/min. During the same procedure, a GRIN lens (Inscopix GLP-0673; 7.3 mm x 0.6 mm) was implanted above the CeA (coordinates from bregma: ML +2.2, AP -1.3, DV 4.5 mm). A GRIN lens was inserted into the brain using a holder (Mightex HLR-GRIN-050) at a rate of approximately 0.1 mm/min, with special care taken to prevent bleeding. After implantation, the sides of a GRIN lens and a skull were covered with surgical glue (Surgibond). Then, dental cement was used to mount a head plate (Neurotar NTR000484- 06) on animals’ heads. The surface of a GRIN lens was covered with a sealant (World Precision Instruments Kwik-Sil). After surgery, mice received painkillers and antibiotics (butorphanol 3 mg/kg, enrofloxacin 5 mg/kg, and tolfenamic acid 3 mg/kg) subcutaneously, for the next 5 days.

### Two-photon microscope

Calcium imaging of CeM neurons was conducted using a custom-built two-photon microscope, based on the project described by Yao P et al. ^60^. The microscope was equipped with a resonant scanning mirror operating at an 8 kHz scanning rate. The signal was collected at 512x512 pixels at 30 Hz, with each frame averaged 3 times. For a two-photon excitation, a water-cooled Halite 920 laser (Fluence) was used. The incorporated laser dispersion management allowed for optimal pulse compression to acquire the best signal-to-noise ratio. We used the 20x Nikon plan-fluor objective with NA = 0.5, which matched the NA of the GRIN lens. Data collection was performed using ScanImage software (mbf BIOSCIENCE).

### Two-photon calcium imaging

Prior to imaging sessions, mice were habituated to head-fixation procedures and exposure to white noise. Before each head-fixation session, mice were lightly anesthetized with isoflurane and head-fixed in a chamber. Habituation was conducted gradually, starting with only 1 minute of head-fixation in darkness. After a week of habituation, imaging sessions started. For five consecutive days, mice were head-fixated, and once the imaging plane on a GRIN lens was identified mice were fully awakened, they were injected subcutaneously with 100 μL of saline or 20 mg/kg cocaine dissolved in saline. Imaging was conducted immediately after the injection. During imaging, a mouse’s face was illuminated with an infrared diode and recorded using an infrared camera (Basler acA2000-165umNIR). The locomotor activity of a mouse was tracked using an encoder that measured rotations of the rotating disc via a Teensy microcontroller (Teensy 3.5 ARM Cortex M4). The infrared camera and an encoder were synchronized with the two-photon microscope, where acquired frames served as a master clock for the other devices.

For most mice, an additional day of imaging was conducted, during which they were exposed to a 7.5% sucrose solution and 0.1 mM quinine delivered through a custom-made lickport.

### Two-photon imaging data analysis

To improve the signal-to-noise ratio, images from a two-photon microscope were averaged two times, which gave an effective frame rate of 5 Hz. Cell segmentation and extraction of the fluorescence signal were performed using Suite2P software ^61^. Subsequently, a custom script written in Matlab was used for the detection of calcium events for individual cells.

Videos from an infrared camera were cropped to the size of a mouse head using FFmpeg software. These cropped videos were then analyzed with DeepLabCut software, which enabled tracking of several points on the mouse head^62^. These tracking points, along with SimBA^63^ software, allowed for the detection of specific mouse movements. Videos of mice face during sugar exposure were analyzed with BehaView software (BehaView 0.0.23 by Dr. Pawel Boguszewski, Laboratory of Behavioral Models, Nencki Institute of Experimental Biology PAS, Warsaw), where licks of sugar solution droplets were manually determined.

### Retrograde labeling of the dopamine-sensitive neurons projecting to the CeA

DRD2-Cre or DRD1-Cre mice were anesthetized with 5% isoflurane and injected i.p. with painkillers (butorphanol 3 mg/kg and tolfenamic acid 3 mg/kg). 50 nl of adeno-associated viruses (pAAV-hSyn-DIO-EGFP at titer 1,5 x 10*8; Addgene 50457-AAVrg or retrograde hSyn-DIO-mCherry at titer 3,75 x 10*8; Addgene 50459-AAVrg) were stereotactically injected into the CeA (coordinates from bregma: ML +2.2, AP -1.3, DV 4.6 mm) at a rate of 10 nl/min. After surgery, the welfare of the mice was monitored over the next 5 days. If needed, mice received painkillers and antibiotics (butorphanol 3 mg/kg, enrofloxacin 5 mg/kg, and tolfenamic acid 3 mg/kg). Three weeks post-surgery mice were perfused with cold PBS followed by 4% PFA dissolved in PBS. Their brains were removed and they were kept in 4% PFA for 24 hours in the 4°C and later in 4°C PBS. Brains were sliced on a vibratome (Leica VT 1200 S) into 50 μm slices. Brain slices were permeabilized at room temperature in 0.25% Triton X-100 in PBS and then incubated for 3 h in a blocking solution (3% BSA and 0.1% Triton X-100 in PBS). Slices were then incubated for 24 hours at 4°C in 0.1% Triton X-100 and 0.1% BSA in PBS containing a mix of antibodies against endogenous fluorophores: anti-GFP (Abcam ab13970, 1:1500) and anti-RFP (ThermoFisher M11217, 1:1500). Later, brain slices were incubated for 3 h at room temperature with a mix of secondary antibodies (ThermoFisher A-11039 or Abcam ab150158, 1:400) that had been previously centrifuged at 20000g for 10 minutes. Brain slices were mounted on coverslips and imaged with the fluorescent microscope (Olympus VS110). Images were then registered on Allen Brain atlas with *Aligning Big Brains* & *Atlases* software (ABBA, <u>BioImaging</u> <u>& Optics Platform</u> at EPFL, https://biop.github.io/ijp-imagetoatlas/).

#### Conditioned place preference

DRD2-Cre or DRD1-Cre mice were anesthetized with 5% isoflurane and injected i.p. with painkillers (butorphanol 3 mg/kg and tolfenamic acid 3 mg/kg). 250 nl of adeno-associated viruses (pAAV-hSyn-DIO-hM4D(Gi)-mCherry^64^ at titer ≥ 7×10¹² vg/mL; Addgene 44362-AAV1 or pAAV-hSyn-DIO-hM3D(Gq)-mCherry^64^ at titer ≥ 7×10¹² vg/mL; Addgene 44361-AAV1) were stereotactically injected into the CeA (coordinates from bregma: ML +2.2, AP -1.3, DV 4.6 mm) at a rate of 75 nl/min. After surgery, the welfare of the mice was monitored over the next 5 days. If needed, mice received a cocktail of drugs (butorphanol 3 mg/kg, enrofloxacin 5 mg/kg, and tolfenamic acid 3 mg/kg). As the “sham” group we included mice that showed no signs of viral infection in any part of the brain, as well as mice that had stereotactically injected 250 nl of saline into the CeA.

Three weeks post-surgery mice underwent a 10-day CPP protocol. Each day, mice were individually placed in the CPP apparatus and their behavior was video recorded for 18 minutes at 25 fps. The CPP apparatus is a cage (82 x 42 cm) divided into three chambers distinguished by wall patterns and floor structure. Chambers are connected by doors that can be either opened to allow free exploration of the entire cage or closed to confine the mouse to a selected chamber. On day 1, all doors were opened and mice were placed in the middle chamber to freely explore the cage to establish their baseline preference. Following days the doors were closed. For the cocaine groups, one of the two side chambers was randomly selected as the drug-paired (“conditioned chamber”) and the other as the vehicle-paired (“non-conditioned chamber”). On days 2, 4, 6, and 8, the cocaine groups received i.p. injections of cocaine hydrochloride (20 mg/kg in saline) and were immediately placed into the drug-paired chamber. On days 3, 5, 7, and 9 animals from cocaine groups received i.p. injections of saline and were immediately put into the vehicle-paired chamber. The saline control groups received only i.p. saline injections daily from day 2 to 9 (in our analysis, the chamber where they received injections on days 2, 4, 6, and 8 was assigned as the “conditioned chamber”, whereas the chamber where they received injections on days 3, 5, 7, 9 was assigned as the “non-conditioned chamber”). On day 10 mice received i.p. injections of Clozapine-N-oxide (1 mg/kg), the ligand for DREADD receptors. 30 minutes later they were put into the middle chamber with all doors open to assess the animals’ preferences.

After the CPP test mice were perfused with PBS followed by 4% PFA dissolved in PBS. Their brains were removed and kept in 4% PFA for 24 hours in the 4°C and later in 4°C PBS. Brains were sliced on a vibratome (Leica VT 1200 S) into 100 μm slices and mounted on coverslips. Viral expression was verified using a fluorescence microscope. Mice without a viral expression were classified as “sham” animals. Video recordings were analyzed with Bonsai software and Python script ^65^.

#### Cocaine and sucrose exposure of freely-moving animals

To study the cocaine’s effect on freely moving, DRD1-TdTomato/DRD2-GFP mice received two 100 μL injections of saline for two days. Then, we injected them with 100 μL of saline for another 7 consecutive days or 100 μL of 20 mg/kg body weight of cocaine hydrochloride dissolved in saline. Immediately after the injection, mice were placed in another, larger cage (33 × 21 cm), and their locomotor activity was recorded for 30 min with a camera suspended above the cage. DeepLabCut software was used to track several points on the mouse body. Based on these tracking points the distance covered by animals was calculated.

For the sweet water exposure experiment, mice were moved to another cage for two hours, where they had access to a bottle with either sweet (7.5% sucrose solution) or fresh water. All mice were habituated to the new cage for two days before the experiment. During that time only freshwater bottle was available. Then, mice were exposed to water or 7.5% sucrose solution for 7 consecutive days. To measure drinking time, we used a previously described device based on the programmable microcontroller Arduino Leonardo, which detects the closing of the electric circuit^6^.

### Statistical analysis

Statistical analysis was conducted using GraphPad Prism 9 software. For each group normality was assessed using the Shapiro-Wilk test, followed by analysis of differences in statistical means. Statistical significance was determined for p < 0.05.

## Supplemental information

A document containing supplementary figures S1-S5, related to Figures 1, 2, 3, 6, and 7, respectively.

## Data and materials availability

All data are available in the main text and the supplementary materials. Confocal images, behavioral data, electrophysiological recordings, calcium activity traces, and facial landmarks tracking are available at the repository https://doi.org/10.18150/TJDGZC

## Notes

### Competing Interest Statement

The authors have declared no competing interest.

