## Supplementary material for "Opposing effects of rewarding and aversive stimuli on D1 and D2 types of dopamine-sensitive neurons in the central amygdala": Suplemental figures 1-5

**This file includes:**

Figs. S1 to S5

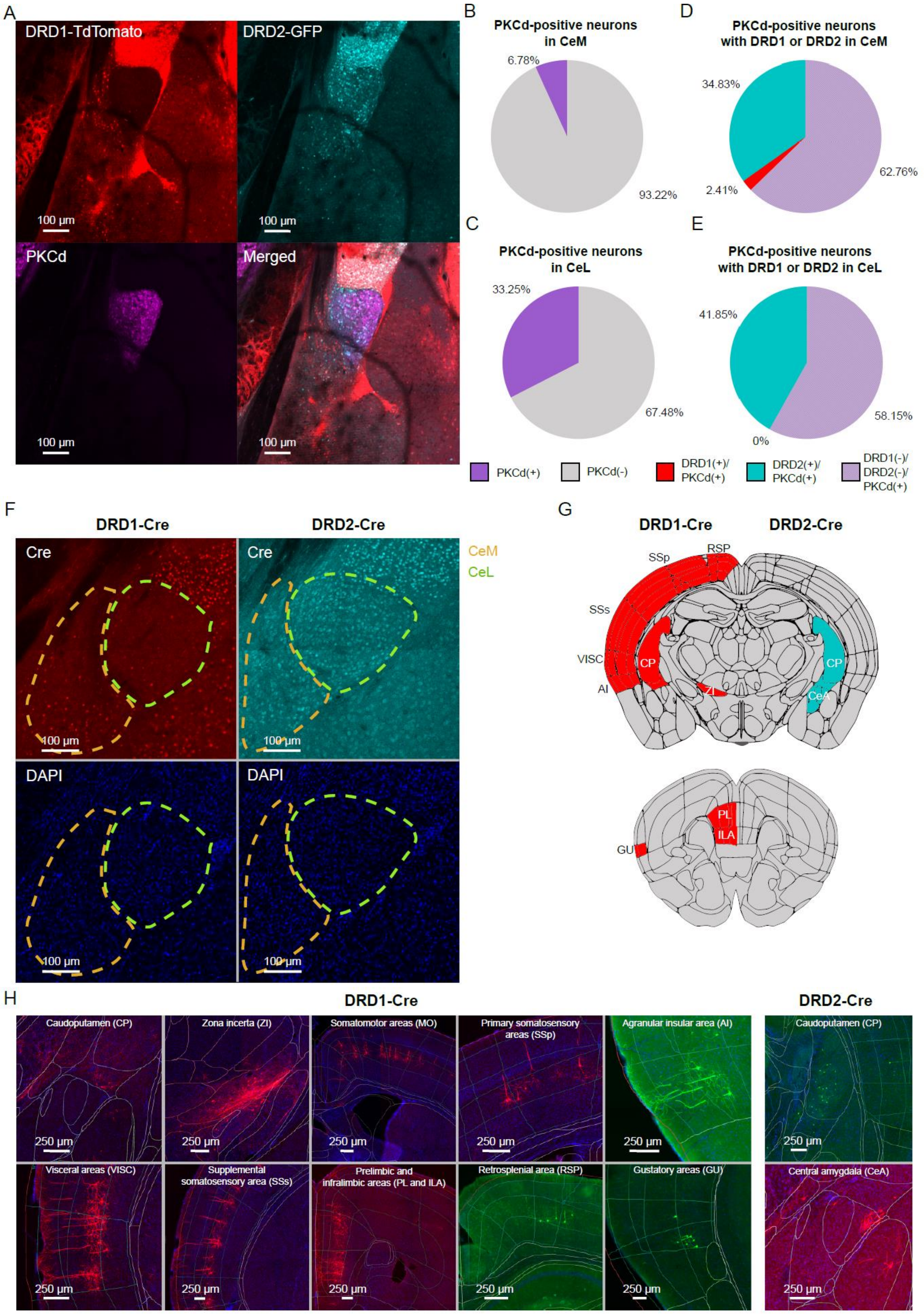
A
DRD1-TdTomato
DRD2-GFP
PKCd
Merged
100 µm
B
PKCd-positive neurons in CeM
6.78%
93.22%
C
PKCd-positive neurons in CeL
33.25%
67.48%
D
PKCd-positive neurons with DRD1 or DRD2 in CeM
34.83%
2.41%
62.76%
E
PKCd-positive neurons with DRD1 or DRD2 in CeL
41.85%
0%
58.15%
PKCd(+)
PKCd(-)
DRD1(+)/ PKCd(+)
DRD2(+)/ PKCd(+)
DRD1(-)/ DRD2(-)/ PKCd(+)
F
DRD1-Cre
DRD2-Cre
Cre
DAPI
CeM
CeL
G
DRD1-Cre
DRD2-Cre
SSp
RSP
SSs
VISC
AI
CP
ZI
CeA
PL
ILA
GU
H
DRD1-Cre
DRD2-Cre
Caudoputamen (CP)
Zona incerta (ZI)
Somatomotor areas (MO)
Primary somatosensory areas (SSp)
Agranular insular area (AI)
Visceral areas (VISC)
Supplemental somatosensory area (SSs)
Prelimbic and infralimbic areas (PL and ILA)
Retrosplenial area (RSP)
Gustatory areas (GU)
Central amygdala (CeA)
250 µm

**Fig. Supp. 1: Markers of the amygdala and projections from dopamine-sensitive cells to the CeA (related to Fig. 1).** A) Co-expression of PKCδ and dopamine receptors in the mouse amygdala. B, C) Percentages of PKCδ neurons among all neurons in the CeM (B) and CeL (C). D, E). PKCδ neurons co-expression with DRD1 and DRD2 in CeM (D) and CeL (E). F) Labelling of Cre protein in brain slices from DRD1-Cre (red) and DRD2-Cre (cyan) mice. Dashed lines represent the borders of CeM (yellow) and CeL (green) nuclei. G) Schematic representation of regions with DRD1(+) (red, left) and DRD2(+) (cyan, right) cells projecting to the CeA. H) Example images of retrograde tracing of DRD1(+) and DRD2(+) neurons projecting to the CeA.
N numbers in groups for CeM: N = 4(15) and for CeL: N = 3(12), where N = number of animals (number of analyzed images).

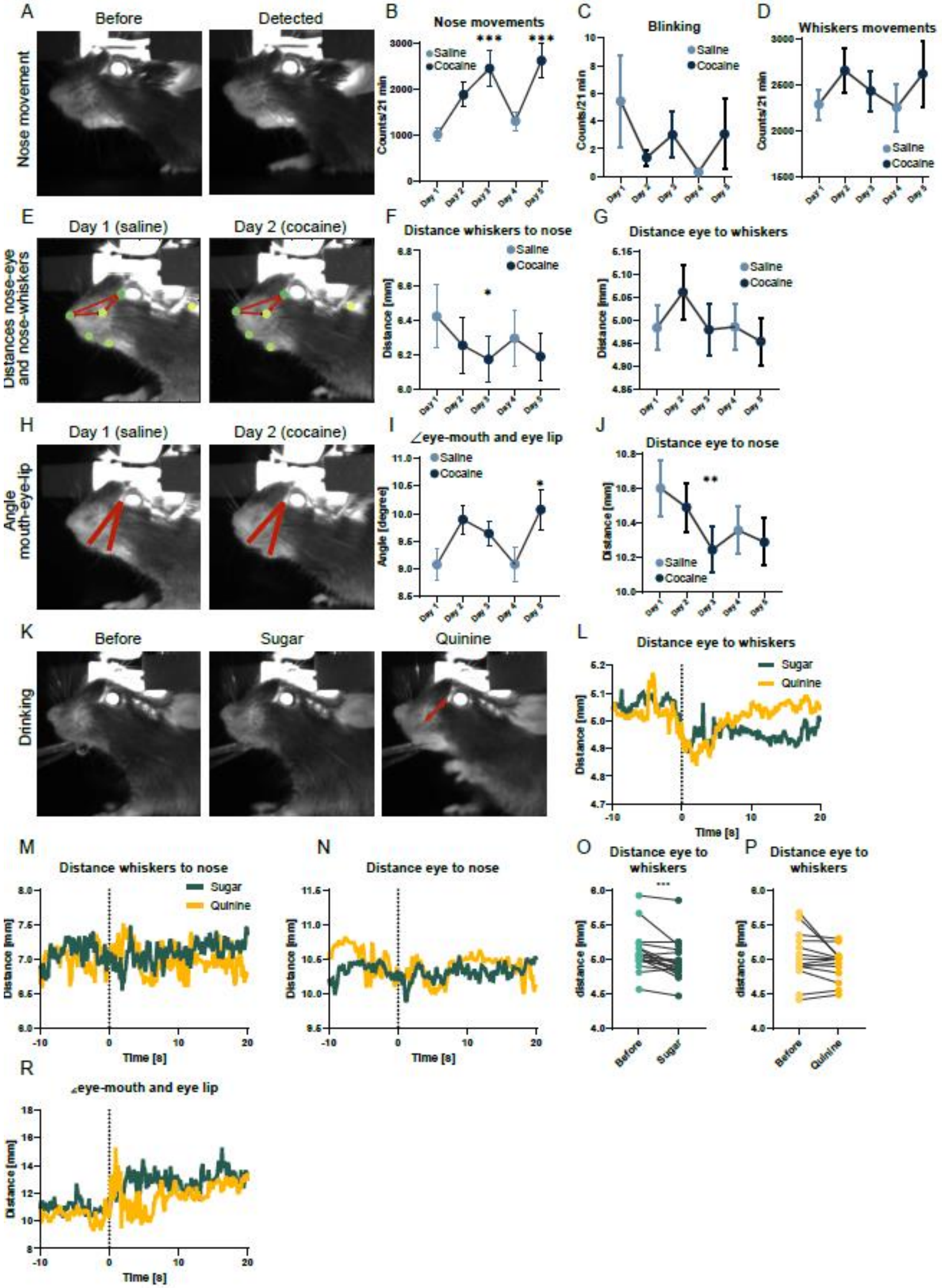
A
Before
Detected
Nose movement
B
Nose movements
Saline
Cocaine
Counts/21 min
C
Blinking
Saline
Cocaine
Counts/21 min
D
Whiskers movements
Saline
Cocaine
Counts/21 min
Day 1
Day 2
Day 3
Day 4
Day 5
E
Day 1 (saline)
Day 2 (cocaine)
Distances nose-eye and nose-whiskers
F
Distance whiskers to nose
Distance [mm]
G
Distance eye to whiskers
Distance [mm]
H
Day 1 (saline)
Day 2 (cocaine)
Angle mouth-eye-lip
I
∠eye-mouth and eye lip
Angle [degree]
J
Distance eye to nose
Distance [mm]
K
Before
Sugar
Quinine
Drinking
L
Distance eye to whiskers
Sugar
Quinine
Distance [mm]
Time [s]
M
Distance whiskers to nose
Sugar
Quinine
Distance [mm]
Time [s]
N
Distance eye to nose
Distance [mm]
Time [s]
O
Distance eye to whiskers
distance [mm]
Before
Sugar
P
Distance eye to whiskers
distance [mm]
Before
Quinine
R
∠eye-mouth and eye lip
Distance [mm]
Time [s]

**Fig. Supp. 2: Facial expressions of mice after cocaine, sugar, and quinine exposure (related to Fig. 2).** A) Example frames from the video recordings before and during nose movement. B, C, D) Graphs representing events counted by SIMBA: nose movements (B), blinking (C), whisker movements (D). Graphs represent averages for all mice ± SEM. E, H) Exemplary frames from a mouse exposed to saline (left) and cocaine (right) showing distances of tracked points (E) and angle between mouth and eye lip (H) Dots represent tracked points of the mouse face and red arrows represent distances between nose, whiskers, and eye. F, G, I, J) Graphs representing distances between whiskers and nose (F), distances between eye and whiskers (G), angles between mouth, lip, and eye (I), and distances between eye and nose during cocaine and saline sessions. Graphs represent averages for all mice ± SEM. K) Example frames of a mouse before and after sugar and quinine droplet lick. Red arrows represent distances between eyes and whiskers. L-N, R) Graph representing averaged distances between eye and whiskers (L), distances between whiskers and nose (M), distances between eye and nose (N), and angles between mouth, eye, and lip (R) 10 seconds before and 20 seconds after a lick of sugar (green) and quinine (yellow). O, P) Graphs representing averaged distances between the eye and whiskers before and after a lick of sucrose solution (O) or quinine solution (P).

N numbers in groups for B-D, F, G, I, and J: N = 17, where N = number of animals. For P-S: N sugar = 20 and N quinine = 15, where N = number of animals. Statistical differences for B-D, F, G, I, and J were measured with one-way ANOVA with Dunnett's multiple comparisons test, with p-values for B day 1 vs. day 3 $p = 0.0029$, and day 1 vs. day 5 $p = 0.0007$, and p-values for F day 1 vs. day 3 $p = 0.0101$, and p values for I day 1 vs. day 5 $p = 0.0278$, and p-values for J day 1 vs. day 3 $p = 0.0035$, day 1 vs. day 4 $p = 0.0017$, and day 1 vs. day 5 $p = 0.0033$. Statistical differences for O and P were measured with the Wilcoxon test, with p-values for O $p = 0.0008$ and P $p = 0.1353$. Statistical differences are represented by stars, where *, **, and *** correspond respectively to $p < 0.05$, $p < 0.01$, and $p < 0.001$.

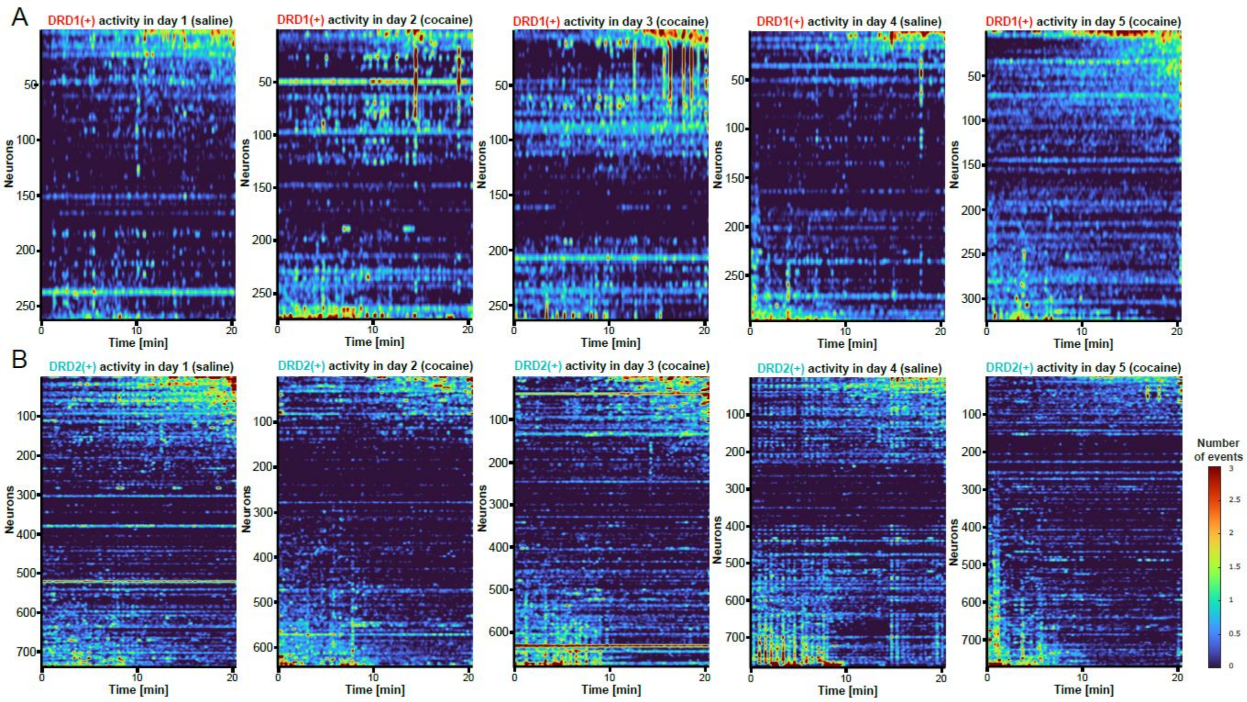


**Fig. Supp. 3: Calcium activity of DRD1- and DRD2-positive neurons in the medial part of the central amygdala after cocaine exposure (related to Fig. 3).** A, B) Heatmaps showing calcium activity of all recorded DRD1(+) (A) and DRD2(+) (B) neurons across the whole experiment. During each session mice received either saline or cocaine injection before imaging. Neurons were sorted based on the difference in their activity before and after 7.5 minutes of recording.

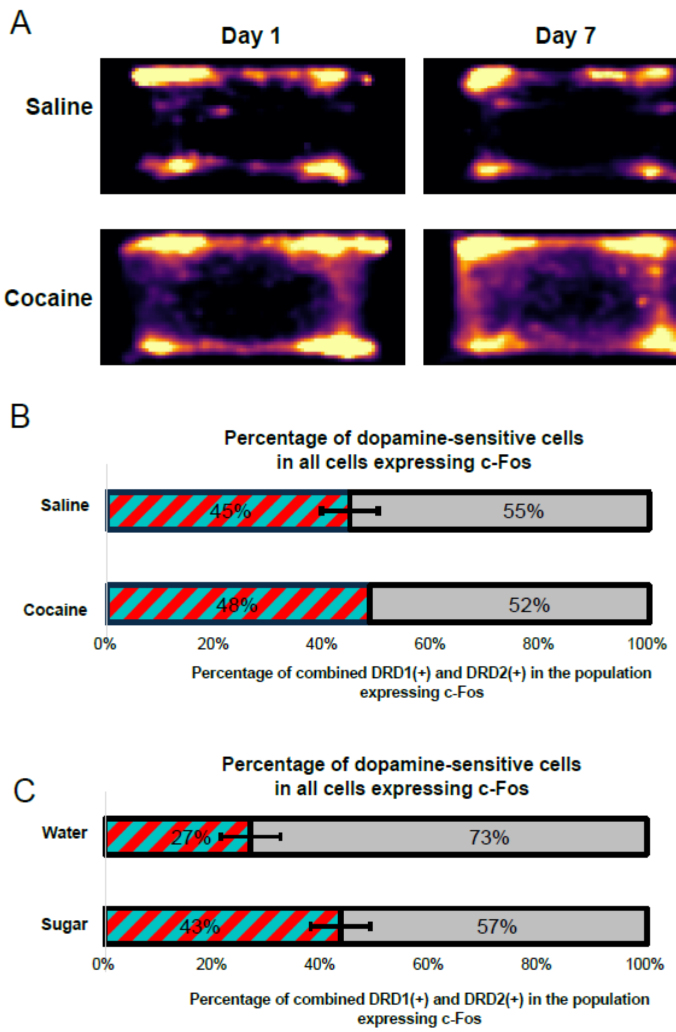


**Fig. Supp 4: Cocaine-induced locomotor activity and c-Fos expression (related to Fig. 6).** A) Heatmaps of the locomotor activity of mice exposed to saline and cocaine after the first cocaine i.p. injection (Day 1) and after 7 injections (1/day; Day 7). B, C) Percentage of dopamine-sensitive cells (striped red and cyan) in the population of all cells expressing c-Fos in the medial nucleus of the central amygdala in mice exposed to sugar (B) or cocaine (C). Graphs represent averages ± SEM.

Number of groups for B and C: N Water = 8(15); N Sugar = 5(11); N Saline = 2(10); N Cocaine = 1(1), where N = number of animals (number of analyzed ROIs).

**Fig. S5.**

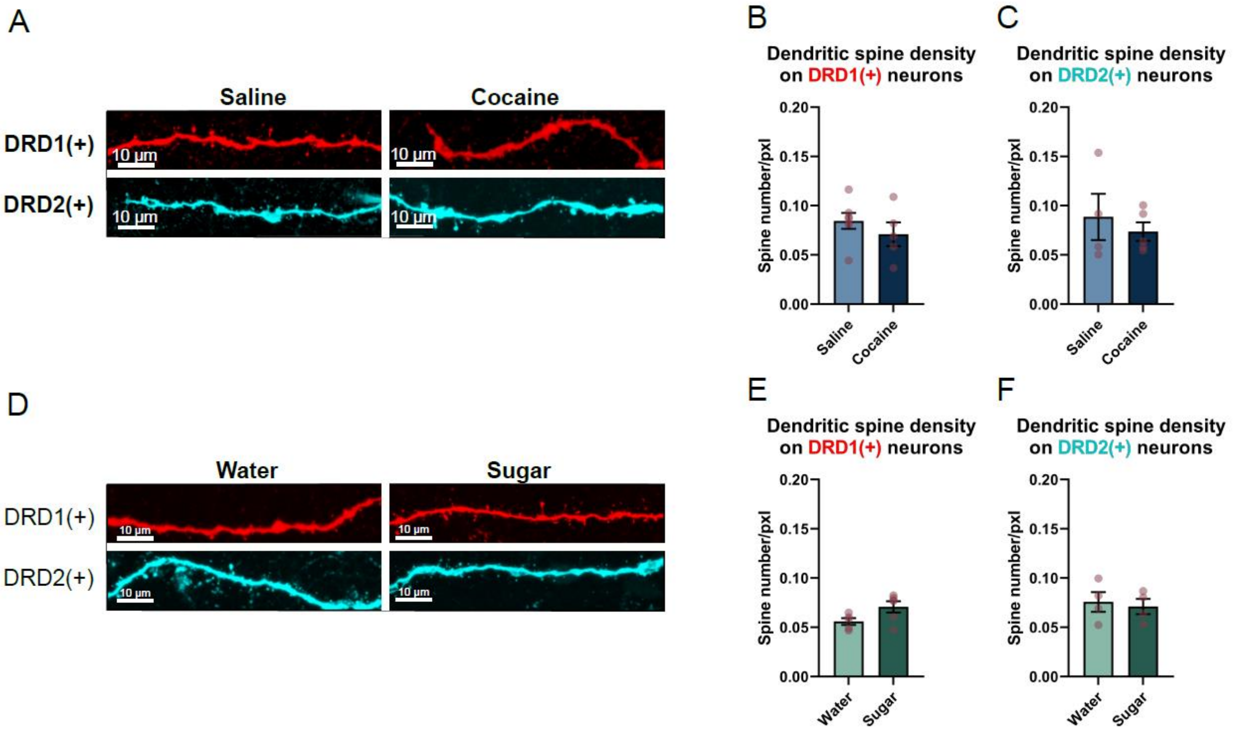


**Fig. Supp. 5: Dendritic spine density of DRD1(+) and DRD2(+) neurons after reward exposure (related to Fig. 7).** A, D) Example images of dendritic spines of DRD1(+) (red) and DRD2(+) (cyan) neurons from brain slices of mice exposed to saline and cocaine (D) or to water and sucrose solution. B, C, E, F) Graphs representing averaged dendritic spine density on DRD1(+) and DRD2(+) neurons from mice exposed to cocaine (B, C) or sucrose solution (E, F). Graphs represent averages ± SEM. Red dots represent values for individual mice.

Graphs represent averages ± SEM. Statistical differences between groups were measured with an unpaired t-test. N numbers in groups for DRD1(+) N Water = 5(39), N Sugar = 6(64), N Saline = 7(89), N Cocaine = 5(17) and for DRD2(+) N Water = 4(19), N Sugar = 4(23), N Saline = 4(23), N Cocaine = 5(33), where N = number of animals (number of analyzed dendrites).
